# PTEN Inhibition Ameliorates Muscle Degeneration and Improves Muscle Function in a Mouse Model of Duchenne Muscular Dystrophy

**DOI:** 10.1101/2020.08.13.249961

**Authors:** Feng Yue, Changyou Song, Di Huang, Naagarajan Narayanan, Jiamin Qiu, Zhihao Jia, Zhengrong Yuan, Stephanie N Oprescu, Bruno T Roseguini, Meng Deng, Shihuan Kuang

## Abstract

Duchenne Muscular Dystrophy (DMD) is caused by mutation of the muscle membrane protein dystrophin and characterized by severe degeneration of myofibers, progressive muscle wasting and loss of mobility, ultimately cardiorespiratory failure and premature death. Here we report that skeletal muscle-specific knockout (KO) of *Phosphatase and tensin homolog* (*Pten*) gene in an animal model of DMD (*mdx* mice) alleviates myofiber degeneration and restores muscle function without increasing tumor incidences. Specifically, *Pten* KO normalizes myofiber size and prevents muscular atrophy, and improves grip strength and exercise performance of *mdx* mice. *Pten* KO also reduces fibrosis and inflammation; and ameliorates muscle pathology in *mdx* mice. Moreover, we found that *Pten* KO upregulates extracellular matrix and basement membrane components positively correlated to wound healing, but suppresses negative regulators of wound healing and lipid biosynthesis; and restores the integrity of muscle basement membrane in *mdx* mice. Importantly, pharmacological inhibition of PTEN similarly ameliorates muscle pathology and improves muscle integrity and function in *mdx* mice. Our finding provides evidence that PTEN inhibition may represent a potential therapeutic strategy to restore muscle function in DMD.

## Introduction

Muscular dystrophy is a group of genetic diseases that cause progressive weakness and loss of muscle mass. Among different types of muscular dystrophy, Duchenne muscular dystrophy (DMD) is the most common type with 1 in about 3,300 live male births affected.^1^ DMD is caused by inherited mutations in the X-linked DMD gene that encodes for the protein dystrophin.^2,3^ Dystrophin, a myofiber membrane protein, is critical for maintaining myofiber stability and integrity,^4–7^ as well as the regenerative capacity of muscle stem cells.^8^ In the absence of dystrophin, DMD patients exhibit severe degeneration of muscle fibers and progressive loss of stem cell-mediated muscle repair.^9^ Unfortunately, restoration of dystrophin expression is extremely challenging due to the enormous size of this protein (427 kDa). In addition to attempts to restore dystrophin expression by gene editing, several therapeutic strategies have been proposed to ameliorate dystrophic phenotypes in animal models by manipulating muscle growth related signaling pathways, including activation of insulin-like growth factor-1 (IGF-1),^10,11^ blockage of the growth-inhibitory myostatin (MSTN) molecule and its receptor activin type IIB (ActRIIB).^12,13^ Furthermore, overexpression of utrophin, a closely related membrane protein with overlapping function with dystrophin, has been examined as a strategy to treat DMD.^14,15^ All these strategies aim to prolong muscle function and life expectancy of DMD patients through preserving or increasing muscle mass. Despite the promising effects in animal models, these strategies are mostly at the preclinical investigation stage and the long-term functional outcomes of these treatments are poor.^16^ Therefore, new strategies are still needed to boost the regrowth of damaged muscle and increase muscle strength in DMD patients, with the ultimate goal to improve their quality of life and survival.

Phosphatase and tensin homolog (PTEN) is a dual-specificity lipid and protein phosphatase that functions through dephosphorylating PIP3 to PIP2 to antagonize PI3K thus inhibiting downstream AKT signaling.^17^ Interestingly, previous studies have reported a marked upregulation of PTEN level in skeletal muscles of DMD patients and animal models, accompanied by muscle weakness and loss.^18–21^ Recent studies have implied that indirect inhibition of Pten signaling by modulating its regulators improves the dystrophic muscle function in animal models of DMD.^18,22,23^ Specifically, skeletal muscle–specific miR-486 overexpression in *DMD^mdx^* animals decreased Pten expression levels, and subsequently increased levels of phosphorylated AKT and improved muscular dystrophy-associated symptoms.^18^ Moreover, repression of phosphatidylinositol transfer protein α (PIPTNA) ameliorates the pathology of DMD by decreasing Pten level.^22^ As miRNA-486 and PIPTNA can potentially affect many downstream targets in addition to Pten, the specific role of Pten in DMD is still unclear.

It has been reported recently that genetic ablation of *Pten* in embryonic myoblasts significantly promotes postnatal muscle growth, induces muscle hypertrophy, improves motor function and protects the mice from denervation-induced muscle atrophy.^24^ However, whether direct Pten inhibition could benefit the dystrophic muscles remains unknown. In the present study, we established *MyoD^Cre^*-driven knockout of *Pten* gene in a clinically relevant *DMD^mdx^* mouse strain to achieve specific inactivation of Pten in myogenic progenitor cells and myofibers. We found that loss of *Pten* ameliorates muscle pathology and preserves muscle mass and function in the *mdx* mice. We also explored the underlying mechanism through which Pten inhibition improves muscle structure and function, and found that loss of *Pten* upregulates extracellular matrix and basement membrane components correlating with MG53 membrane repair signaling to restore the integrity of muscle basement membrane. We further demonstrate that pharmacological inhibition of Pten remarkably ameliorates the symptoms of *mdx* mice and improves the dystrophic muscle function. These results indicate that direct targeting of Pten represents a potential therapeutic strategy for treatment of DMD.

## Results

### Myogenic *Pten* Knockout Prevents Muscle Wasting in *mdx* Mice

We analyzed two publicly available microarray datasets on skeletal muscle biopsies from 23 DMD patients and 22 unaffected control patients,^21,25^ and found that *PTEN* mRNA level was significantly upregulated in the skeletal muscles of DMD patients (Figures S1A and S1B). We also examined the protein level of Pten in both fast and slow contracting muscles of *mdx* mice. The results showed that Pten level is increased mildly in the fast extensor digitorum longus (EDL) (Figures S1C and S1D), and significantly increased in slow soleus muscles of the mdx relative to corresponding muscles in WT mice (Figures S1E and S1F). To further examine if upregulation of Pten is due to muscle cells or non-muscle cells (such as fibroblasts) in the muscle tissues, we examined the Pten level in enzymatically dissociated EDL myofibers free of non-muscle cells. Again, Pten protein levels were much higher in EDL myofibers of mdx mice than in those of WT mice (Figures S1G and S1H). These results confirm that Pten level is elevated in dystrophic muscles of DMD patients and mdx mice.

To directly study the role of PTEN in DMD, we established the *MyoD^Cre^*:*Pten^f/f^*:*DMD^mdx^* (*mdx/Pten^cKO^*) mouse model to achieve the specific deletion of *Pten* in myogenic progenitor cells and myofibers. We used the newly established D2.B10 *DMD^mdx^* (D2-*mdx*) mouse strain, which better recapitulates the characteristics of DMD myopathology including earlier onset of pathology (before 7 weeks old), lower hind limb muscle weight, increased fibrosis and remarkable muscle weakness.^26–28^ We included littermate *Pten^f/f^* mice with D2.B10 background as control for most of the experiments, and syngenic C57BL/6J mice of identical age and sex as control for part of the study to better assess muscle phenotypes and functions of *mdx* mice in the absence of *Pten*. All these control mice were named wild-type (WT) in this study. A hallmark of DMD is the progressive muscle loss. We first evaluated if *Pten* KO could preserve muscle mass in *mdx* mice. Morphologically, the *mdx/Pten^cKO^* mice were larger than the littermate *mdx* mice, with 11.2% increase in body weight at 7 weeks of age (Figures 1A and 1B). Body composition measurement by EchoMRI showed that *mdx* mice had lower lean mass and higher fat mass compare to WT mice, while *Pten* KO in *mdx* mice normalized the body weight and lean mass to similar levels as those in the WT mice, and lowered the fat mass even below the WT level (Figures 1C and 1D). Additionally, we did not observed any increased incidence of spontaneous rhabdomyosarcoma-like tumors in both young (~3 months, n ≥ 30 for each genotype) and old (~12 months, n ≥ 7 for each genotype) adult *mdx* and *mdx/Pten^cKO^* mice.

**Figure 1.**
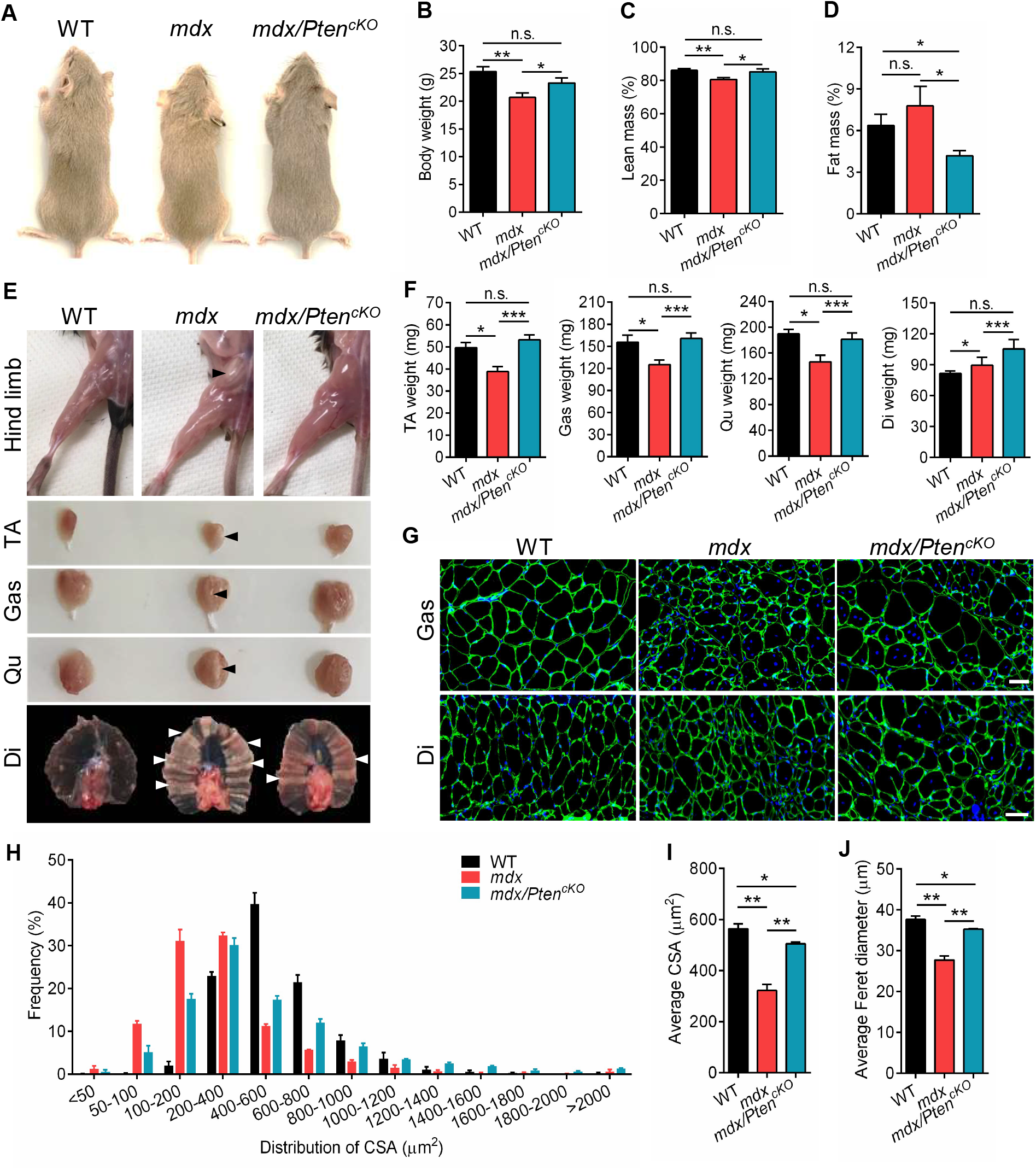
*Pten* Deletion Prevents Muscle Wasting in *mdx* Mice. (A) Representative image of the whole body of 7-week old WT, mdx and *mdx/Pten^cKO^* mice. (B) Body weight of WT, *mdx* and *mdx/Pten^cKO^* mice at 9 weeks of age. n=8 mice (WT), n=15 mice (*mdx*), n=14 mice (*mdx/Pten^cKO^*). (C and D) Percentage of lean mass (C) and fat mass (D) measured by EchoMRI for WT, *mdx* and *mdx/Pten^cKO^* mice at 9 weeks of age. n=6 mice (WT and *mdx*), n=5 mice (*mdx/Pten^cKO^*). (E) Representative image of hind limb, TA, Gas, Qu and Di muscles of WT, *mdx* and *mdx/Pten^cKO^* mice at 9 weeks of age. Arrows indicates fibrosis in muscles. (F) Muscle weight of WT, *mdx* and *mdx/Pten^cKO^* mice at 9 weeks of age. For TA, Gas and Qu muscles, n=8 mice (WT), n=15 mice (*mdx* and *mdx/Pten^cKO^*). For Di muscles, n=5 mice (WT, *mdx* and *mdx/Pten^cKO^*). (G) Representative immunofluorescence staining of α-Laminin for Gas and Di muscles in WT, *mdx* and *mdx/Pten^cKO^* mice. Scale bar: 50 μm. (H) The frequency of cross-sectional area (CSA) distribution for Di muscles in WT, *mdx* and *mdx/Pten^cKO^* mice at 9 weeks of age. n=3 mice (WT, *mdx* and *mdx/Pten^cKO^*). Average 2400 myofibers were measured for each group. (I) The average CSA of Di muscles in WT, *mdx* and *mdx/Pten^cKO^* mice. n=3 mice (WT, *mdx* and *mdx/Pten^cKO^*). (J) The average Feret diameters of Di myofbers in WT, *mdx* and *mdx/Pten^cKO^* mice. n=3 mice (WT, *mdx* and *mdx/Pten^cKO^*). Data are means ± s.e.m. Two-tailed *t*-test: \**P* < 0.05; \*\**P* < 0.01; \*\*\**P* < 0.001; n.s., *P* > 0.05.

To determine if the muscle mass change contributes to the increased body weight and lean mass, different hind limb muscles were examined after euthanasia. The hind limb muscles including tibialis anterior (TA), gastrocnemius (Gas) and quadriceps (Qu) muscles of adult *mdx* mice were all significantly smaller than those of age-matched WT mice at 9 weeks of age (Figure 1E), indicating muscle wasting in the *mdx* mice. Compared to *mdx* mice, *mdx/Pten^cKO^* increased the weights of TA, Gas, Qu and diaphragm (Di) muscles by 37%, 28%, 24% and 18%, respectively, to similar levels of WT mice (Figure 1F). Notably, the Di muscle weight of *mdx/Pten^cKO^* mice was even higher than that of the WT mice (Figure 1F). Histologically, the Gas and Di myofibers of *mdx/Pten^cKO^* mice appeared larger than those of *mdx* mice (Figure 1G). The distribution curve of cross-sectional areas (CSA) of Gas and Di myofibers of *mdx/Pten^cKO^* mice showed a clear right shift when overlaid to that of *mdx* mice (Figures 1H and S2A), resulting in a significant increase of average CSA (Figures 1I and S2B), and Feret diameters (Figures 1J and S2C) to similar levels observed in the WT mice. These results demonstrate that *Pten* KO preserves the muscle mass that is otherwise wasted in the *mdx* mice.

### Loss of *Pten* Improves Dystrophic Muscle Function

The main symptom of DMD is progressive muscle weakness which eventually leads to immobility and respiratory failure.^29^ To examine if inhibition of Pten improves muscle strength and motor function in *mdx* mice, we measured the whole limb grip strength in 8-week old mice. The grip strength in *mdx* mice (0.82 N) was significantly weaker than that of WT mice (2.15 N), while *Pten* KO significantly improved the muscle grip strength of *mdx* mice by 41 % (Figure 2A). We further subjected these mice to treadmill test to evaluate their exercise performance. The maximum running speed, total running time and total running distance of *mdx* mice was significantly decreased when compared to WT mice (Figures 2B–D). Strikingly, *Pten* KO led to 19 %, 28 % and 41 % increase in maximum running speed, total running time and running distance, respectively, to levels comparable to those of WT mice (Figures 2B–D).

**Figure 2.**
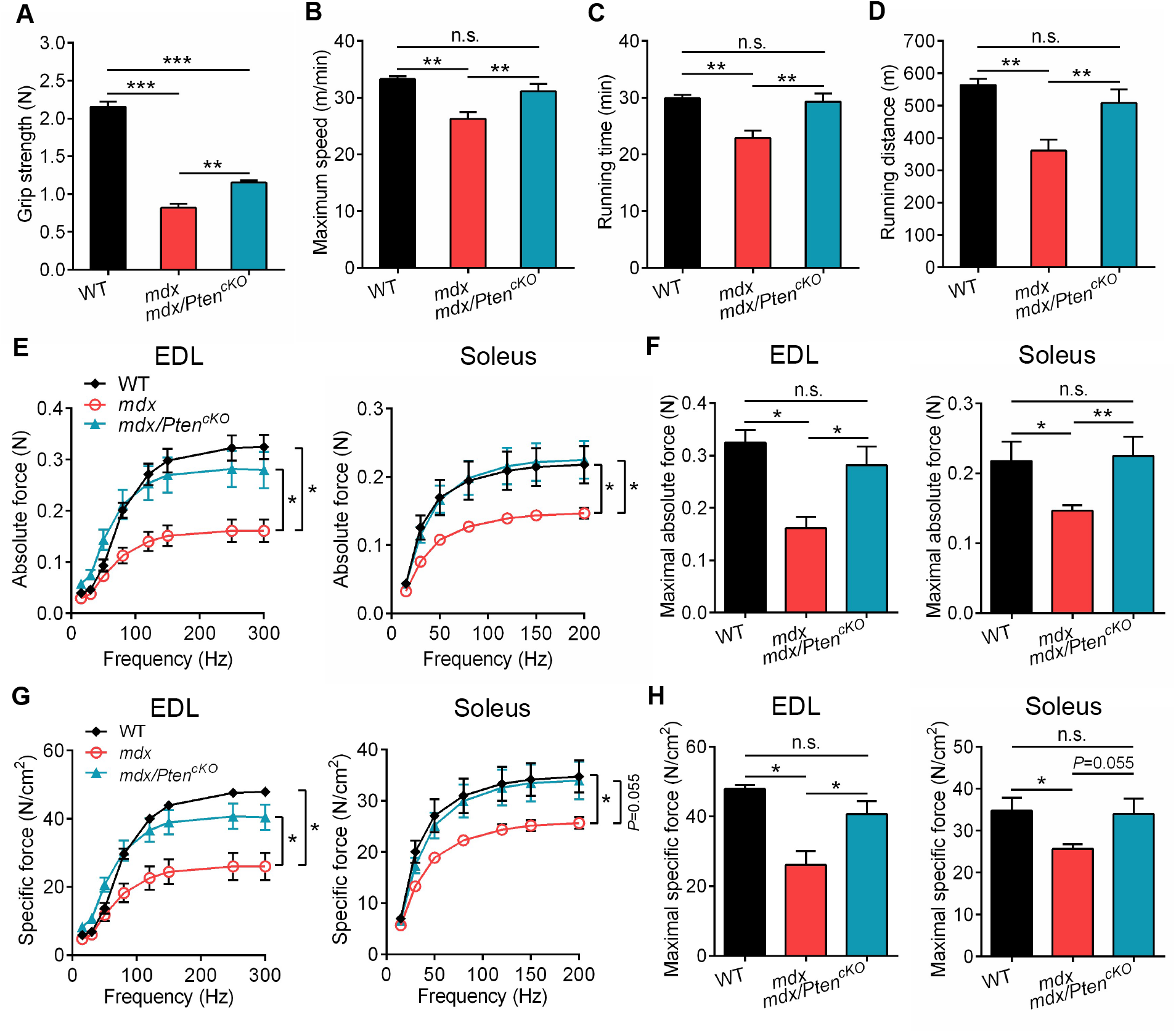
*Pten* Deletion Improves Muscle Motor Function in *mdx* Mice. (A) Gripping strength test of WT, *mdx* and *mdx/Pten^cKO^* mice at 8 weeks of age. n=6 mice (WT), n=8 mice (*mdx* and *mdx/Pten^cKO^*). (B-D) Maximum speed (B), total running time (C), and running distance (D) of WT, *mdx* and *mdx/Pten^cKO^* mice measured by treadmill at 8 weeks of age. n=4 mice (WT), n=7 mice (*mdx* and *mdx/Pten^cKO^*). (E) Absolute force of EDL and Sol muscles from WT, *mdx* and *mdx/Pten^cKO^* mice at 7 weeks of age. For EDL muscle, n=3 mice (WT), n=4 mice (*mdx* and *mdx/Pten^cKO^*); For Sol muscle, n=3 mice (WT and *mdx/Pten^cKO^*), n=4 mice (*mdx*). (F) Maximal absolute force of EDL and Sol muscles from WT, *mdx* and *mdx/Pten^cKO^* mice at 7 weeks of age. n=3 mice (WT, *mdx* and *mdx/Pten^cKO^*). (G) Specific force of EDL and Sol muscles from WT, *mdx* and *mdx/Pten^cKO^* mice at 7 weeks of age. For EDL muscle, n=3 mice (WT), n=4 mice (*mdx* and *mdx/Pten^cKO^*); For Sol muscle, n=3 mice (WT and *mdx/Pten^cKO^*), n=4 mice (*mdx*). (H) Maximal specific force of EDL and Sol muscles from WT, *mdx* and *mdx/Pten^cKO^* mice at 7 weeks of age. n=3 mice (WT, *mdx* and *mdx/Pten^cKO^*). Data are means ± s.e.m. Two-tailed *t*-test: \**P* < 0.05; \*\**P* < 0.01; \*\*\**P* < 0.001; n.s., *P* > 0.05.

To determine if the improvements of grip strength and motor function in *mdx/Pten^cKO^* mice are specifically due to muscle functional improvements, we further evaluated the contractile function of the (fast) EDL and (slow) soleus muscles in WT, *mdx* and *mdx/Pten^cKO^* mice. As expected, the absolute contractile forces responded dose-dependently to varying frequencies of electric stimulation, reaching plateaus at around 250 Hz and 150 Hz in the fast contracting EDL and slow contracting soleus muscles, respectively, regardless of genotypes (Figure 2E). Also consistent with the contractile properties of fast and slow muscles, the maximal contractile force of EDL (~0.32 N) was significantly higher than that of the soleus (~0.22 N) in the WT mice (Figure 2F). Importantly, the dose response kinetics and maximal contractile force of the *mdx/Pten^cKO^* EDL and SOL muscles were indistinguishable from those of the WT mice. In contrast, the absolute contractile forces of *mdx* muscles were significantly lower at all the frequencies examined (Figures 2E and 2F). Specifically, the maximal absolute force of EDL and soleus was increased by 75 % and 53 % in *mdx/Pten^cKO^* mice compared to *mdx* mice (0.28 N versus 0.16 N for EDL, 0.23 N versus 0.15 N for soleus, Figures 2E and 2F). Identical trends were observed when the absolute contractile forces are converted to specific force by taking into consideration of muscle cross-sectional area (Figure 2G). The maximal specific force of EDL and soleus was increased by 56 % and 32 % in *mdx/Pten^cKO^* mice compared to *mdx* mice (40.7 versus 26.1 N/cm^2^ for EDL, 34 versus 25.7 N/cm^2^ for soleus, Figure 2H). These results provide compelling evidence that *Pten* KO rescued the defect of muscle contractile function in *mdx* mice.

### *Pten* Deletion Ameliorates Muscle Pathology in *mdx* Mice

DMD is characterized by continuous degeneration, necrosis, inflammatory infiltrates, and fibrosis.^29,30^ We hence investigated whether *Pten* KO in muscle cells could ameliorate these pathological indices of *mdx* mice. The overall histopathology of the Gas, Qu and Di muscles was assessed in hematoxylin and eosin (H&E) staining sections. We observed severe necrosis and fibrosis in muscles of *mdx* mice, compared to the rare conditions in the WT (Figures 3A, 3B and S3A). Notably, the overall necrotic and fibrotic areas were reduced in the *mdx/Pten^cKO^* compared to the *mdx* mice (Figures 3A, 3B and S3A). We further quantified the necrotic area in these muscles after immunostaining of α-Laminin and DAPI (Figures 3C and S3B). The necrotic areas (indicated by high DAPI density) occupied 55%, 46% and 23% of the Di, Qu and Gas muscle cross-section in *mdx* mice, whereas these were significantly reduced to 25%, 20% and 16% in *mdx/Pten^cKO^* mice, respectively (Figures 3D, 3E and S3C). Masson’s trichrome staining was also used to visualize the fibrosis in muscle by detecting collagen fibers (Figures 3F and S3D). The fibrotic areas in Di and Gas muscles of *mdx* mice were 53% and 42%, in comparison of 2.3% and 0.4% in WT mice, respectively (Figures 3G and S3E). *Pten* KO significantly reduced the collagen deposition by 79% and 81% in Di and Gas muscles of *mdx* mice, respectively (Figures 3G and S3E). In addition, we examined the macrophage infiltration by CD68 staining (Figure 3H). The number of CD68^+^ cells in the Di muscle of *mdx* mice was 35-fold of that in the WT muscles, but the number was significantly decreased in *mdx/Pten^cKO^* muscles relative to *mdx* muscles (18 versus 46 cells per area, Figure 3I). Thus, *Pten* KO leads to significant improvements in various pathological indices in the *mdx* mice.

**Figure 3.**
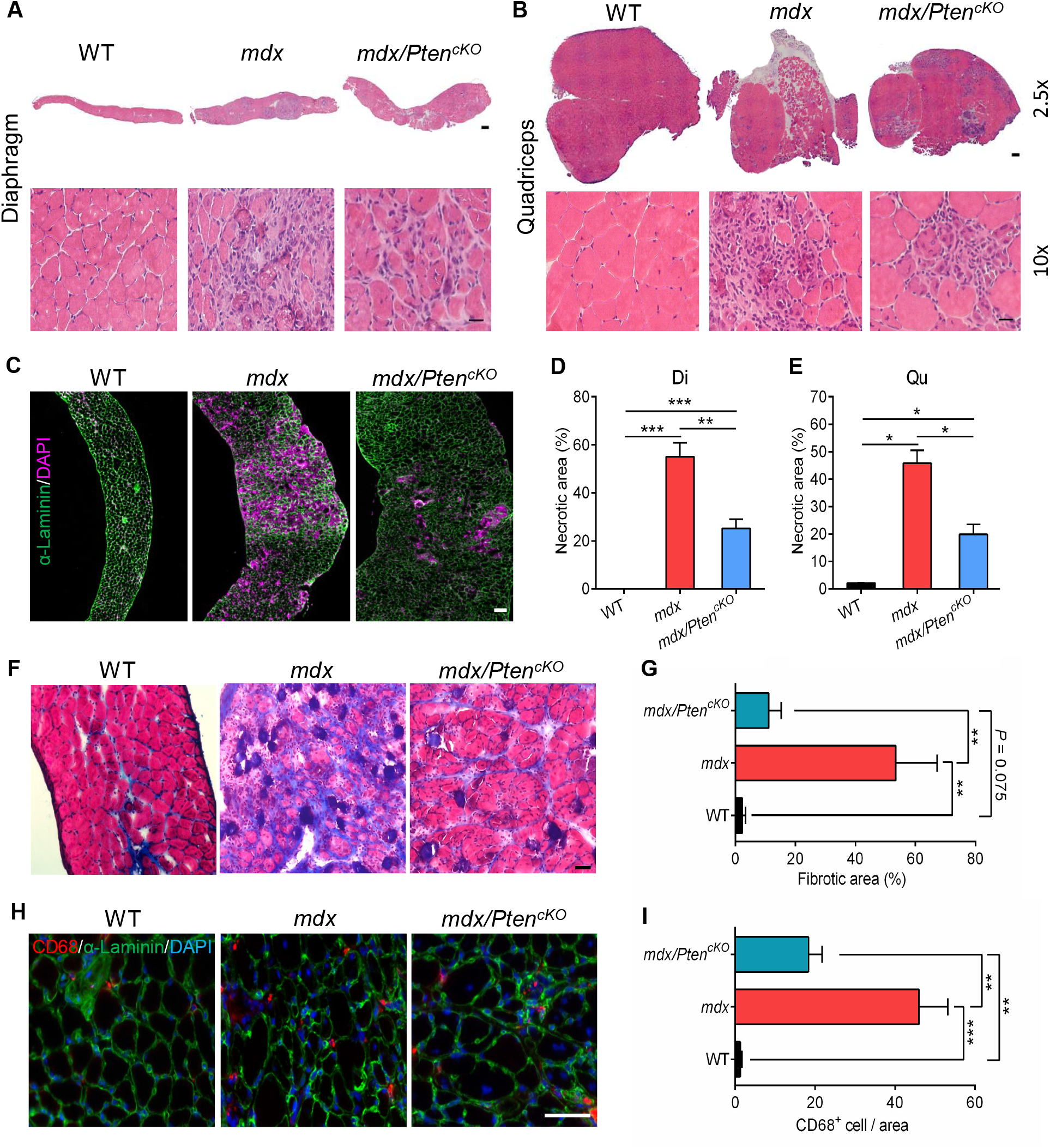
Loss of *Pten* Reduces Muscle Pathology of *mdx* Mice. (A and B) Representative images for HE staining of Qu (A) and Di (B) muscles of WT, *mdx* and *mdx/Pten^cKO^* mice at 9 weeks of age. Scale bar: 500 μm for (A) and 200 μm for (B) at 2.5x, 50 μm at 10x. (C) Representative images of α-Laminin and DAPI staining showing the necrotic area on cross-section of Di muscles. Scale bar: 100 μm. (D and E) Quantification analysis of necrotic area in Qu (D) and Di (E) muscles of WT, *mdx* and *mdx/Pten^cKO^* mice. n=3 mice (WT, *mdx* and *mdx/Pten^cKO^*). (F) Representative images of Masson’s trichrome staining on cross-section of Di muscles from WT, *mdx* and *mdx/Pten^cKO^* mice. Scale bar: 50 μm. (G) Quantification analysis of the percentage of fibrotic area in Di muscles from WT, *mdx* and *mdx/Pten^cKO^* mice. n=4 mice (WT), n=6 mice (*mdx* and *mdx/Pten^cKO^*). (H) Representative images of CD68^+^ and α-Laminin staining on cross-section of Di muscles from WT, *mdx* and *mdx/Pten^cKO^* mice. Scale bar: 50 μm. (I) Quantification analysis of the percentage of CD68^+^ cells in Di muscles from WT, *mdx* and *mdx/Pten^cKO^* mice. n=4 mice (WT), n=5 mice (*mdx* and *mdx/Pten^cKO^*). Data are means ± s.e.m. Two-tailed *t*-test: \**P* < 0.05; \*\**P* < 0.01; \*\*\**P* < 0.001; n.s., *P* > 0.05.

### *Pten* KO Improves Membrane Integrity of *mdx* Myofibers

The etiology of DMD can be explained by membrane leakage due to the lack of dystrophin that resulted in degeneration of myofibers.^29^ To determine whether *Pten* KO affects the membrane integrity, we performed Evans blue (EB) uptake assay in which intraperitoneally injected EB penetrates non-specifically into any cells with leaky membranes. At 24 h after injection, the EB dye accumulated abundantly in all hind limb muscles of *mdx* mice but with very little accumulation in WT muscles (Figure 4A), confirming that loss of dystrophin increases membrane permeability. Notably, *Pten* KO in the *mdx* background significantly blunted EB dye uptake into the muscles of *mdx/Pten^cKO^* mice. A closer examination of isolated Gas, Qu and Di muscles indicated that the WT muscles were free of any EB uptake, and the *mdx/Pten^cKO^* muscles had much lighter EB signal than those the *mdx* muscles (Figure 4A). This observation was further confirmed by fluorescent imaging of EB on cross-sections of Di and Qu muscles (Figures 4B and S4A). The percentage of EB staining area was 22.5-fold and 2.8-fold lower in Di and Qu muscles in *mdx/Pten^cKO^* than in *mdx* muscles, respectively (Figures 4C and S4B). Moreover, we examined the creatine kinase (CK) level in serum, which is a clinical hallmark of the DMD disease as it is released from the injured muscle. Consistent with our EB uptake assay, we found that serum CK level was increased remarkably by 9.4-fold in *mdx* mice compared to WT mice. Strikingly, loss of *Pten* significantly reduced the serum CK level in *mdx* mice by 51% (Figure 4D). Thus, *Pten* KO significantly reduced the leakiness of dystrophic myofibers in *mdx* mice.

**Figure 4.**
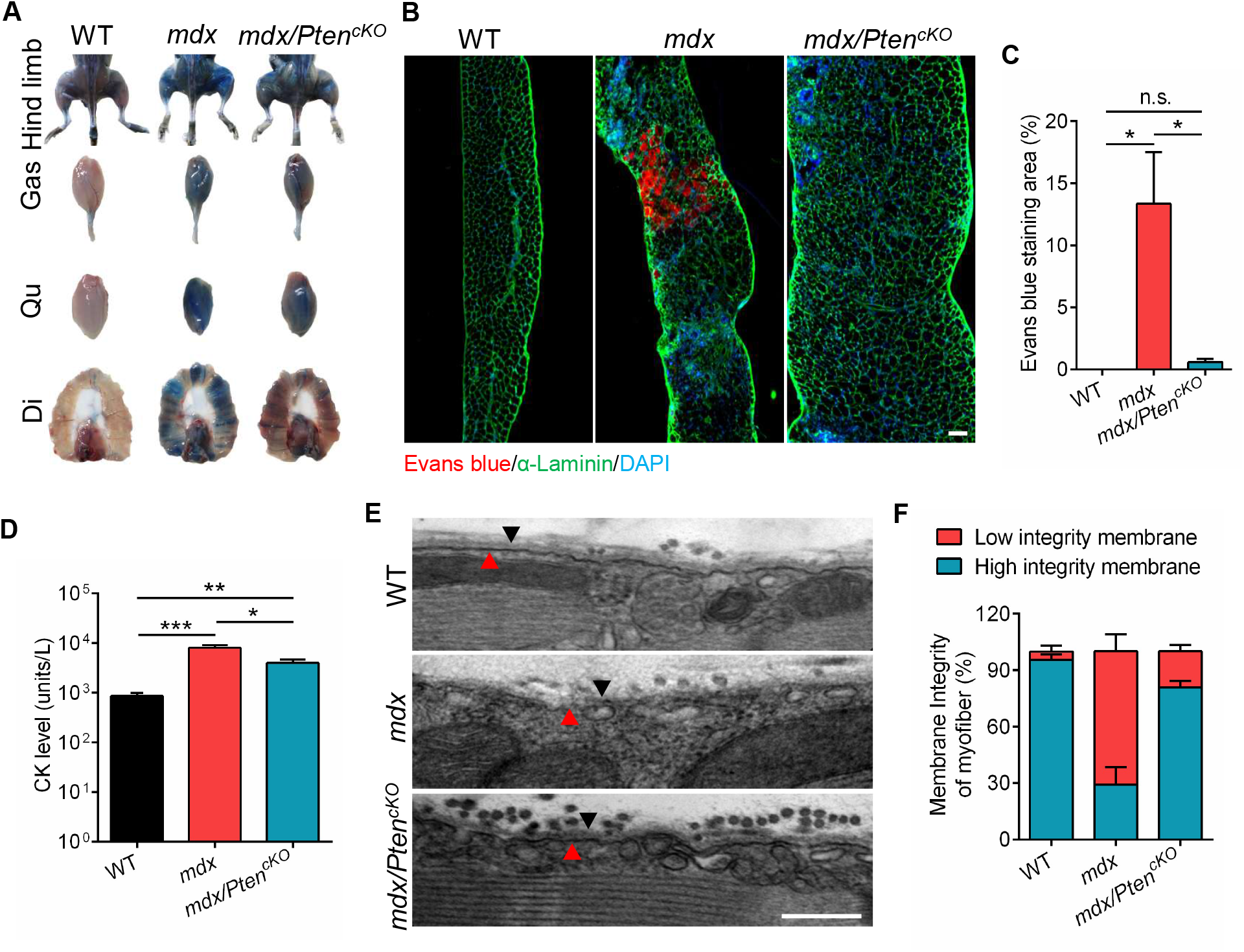
*Pten* Inhibition Protects Against Muscle Damage. (A) Representative images of Evans blue accumulation in hind limb, Gas, Qu and Di muscles of WT, *mdx* and *mdx/Pten^cKO^* mice at 9 weeks of age. (B) Representative images of Evans blue staining on cross-section of Di muscles from WT, *mdx* and *mdx/Pten^cKO^* mice. Scale bar: 100 μm. (C) Quantification analysis of Evans blue staining area on cross-section of Di muscles. n=4 mice (WT, *mdx* and *mdx/Pten^cKO^*). (D) Measurement of CK activity of the serum from WT, *mdx* and *mdx/Pten^cKO^* mice. n=5 mice (WT, *mdx* and *mdx/Pten^cKO^*). (E) Representative TEM images of EDL muscle longitudinal-sections from WT, *mdx* and *mdx/Pten^cKO^* mice. Black arrows indicate the extracellular matrix layer, red arrows indicate sarcolemma layer. Scale bar: 250 nm. (F) Quantification analysis of the percentage of membrane integrity based on TEM images of EDL muscles from WT, *mdx* and *mdx/Pten^cKO^* mice. n=3 mice (WT, *mdx* and *mdx/Pten^cKO^*), 20 myofibers were imaged for each mouse under TEM. Data are means ± s.e.m. Two-tailed *t*-test: \**P* < 0.05; \*\**P* < 0.01; \*\*\**P* < 0.001; n.s., *P* > 0.05.

We further subjected muscles to transmission electron microscopy (TEM) to examine the ultrastructure of the sarcolemma in muscles of WT, *mdx* and *mdx/Pten^cKO^* mice. Broken myofibers were readily observed on the longitudinal-sections of EDL muscle in *mdx* mice (Figure S4C). In contrast, such broken myofibers were observed rarely in EDL muscles of WT and *mdx/Pten^cKO^* mice (Figure S4C). A closer examination revealed that the sarcolemma of EDL and Qu muscles in WT mice appeared as a dark line that was continuous without obvious breakage, whereas the sarcolemma of *mdx* myofibers were fussy and non-continuous (Figures 4E and S4D, red arrows). Notably, *Pten* KO restored the integrity of the sarcolemma in *mdx/Pten^cKO^* muscles (Figures 4E and S4D, red arrow in the bottom panel). Moreover, while the extracellular matrix (ECM) appeared as a clear layer in both WT and *mdx/Pten^cKO^* myofibers, the ECM of *mdx* muscles was hardly recognizable (Figures 4E and S4D, black arrows). The level of membrane integrity in myofibers was then quantified by the presence or absence of clear sarcolemma and ECM layers. We found that the percentage of high integrity membrane increased from 29% in EDL of *mdx* mice to 81% in *mdx/Pten^cKO^* mice, respectively (Figure 4F). These results together demonstrate that *Pten* deletion improves muscle membrane integrity of *mdx* myofibers.

### *mdx/Pten^cKO^* Muscles Showed Gene Enrichments Positively toward Membrane Integrity and Negatively toward Coagulation and Immune Response

To explore the mechanism underlying the improvements of muscle function induced by *Pten* KO, we performed RNA-sequencing analyses comparing transcriptomes of the Di muscles from *mdx* and *mdx/Pten^cKO^* mice. As displayed by volcano diagram, among the total 1,362 differentially expressed genes (adjusted *P* value <0.05), 1,259 genes were downregulated, and 103 genes were upregulated in *mdx/Pten^cKO^* relative to *mdx* muscles (Figure 5A, Data S1 and S2), suggesting that *Pten* KO mainly represses gene expression. Heatmap Cluster Analysis nicely distinguished the expression pattern of differentially expressed genes in *mdx* and *mdx/Pten^cKO^* muscles (Figure 5A).

**Figure 5.**
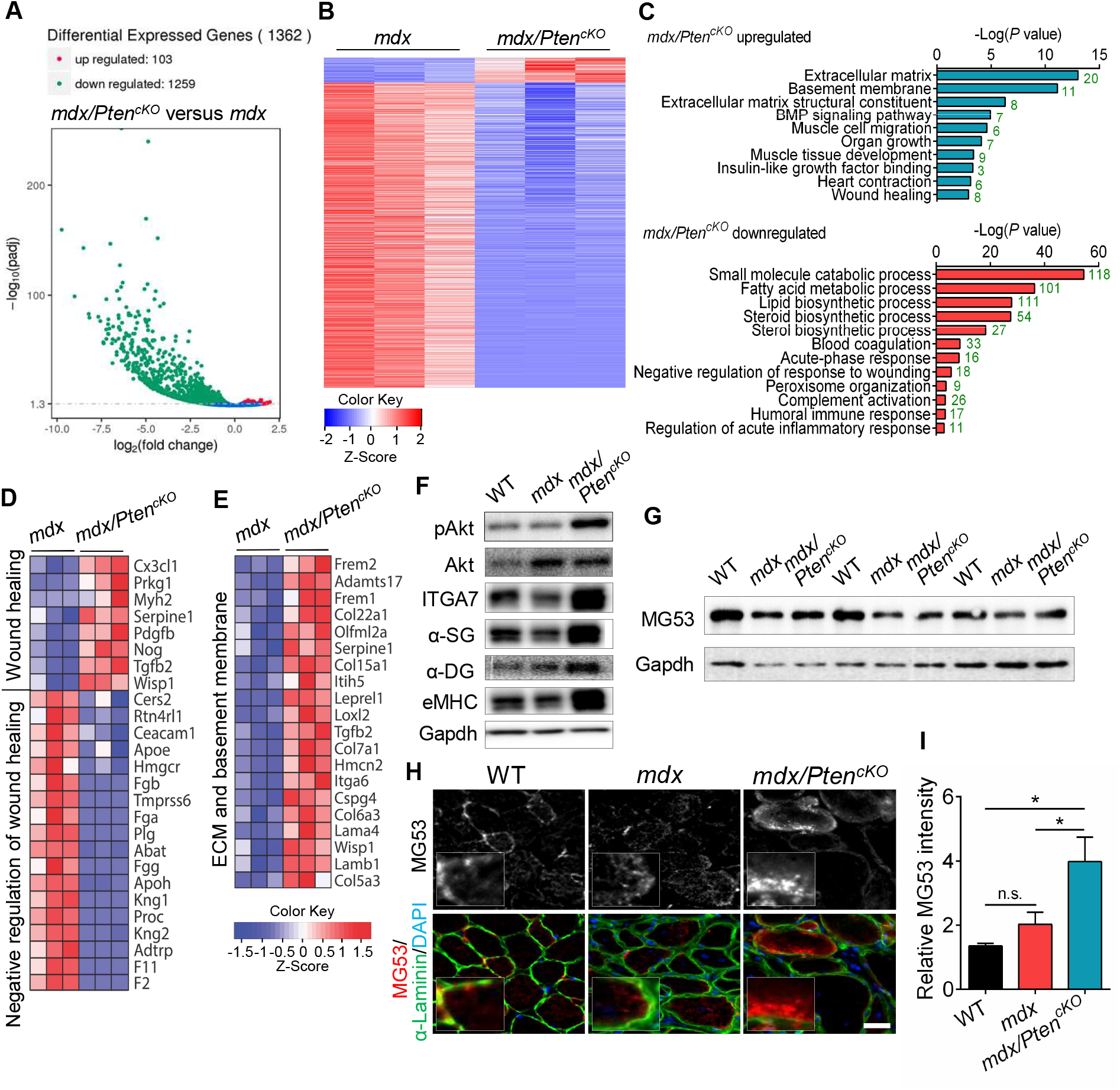
Upregulation of Membrane Repair Integrity Pathways in *mdx* Muscle with *Pten* KO. (A) Volcano diagram of differentially expressed genes in Di muscle of *mdx/Pten^cKO^* versus *mdx* mice. The point represents gene, blue dots indicate no significant difference in genes, red dots indicate upregulated differential expression genes, and green dots indicate downregulated differential expression genes. n=3 mice (*mdx* and *mdx/Pten^cKO^*). (B) Heat map diagram showing the cluster analysis of differentially expressed genes in Di muscles of *mdx* and *mdx/Pten^cKO^* mice. n=3 mice (*mdx* and *mdx/Pten^cKO^*). (C) Gene Ontology analyses showing the enrichment of functional categories in Di muscle of *mdx/Pten^cKO^* versus *mdx* mice. n=3 mice (*mdx* and *mdx/Pten^cKO^*). The number in green color indicates the gene number enriched in the category. (D) Heat map diagram showing the upregulated genes related to wound healing and downregulated genes related to negative regulation of wound healing in Di muscle of *mdx/Pten^cKO^* versus *mdx* mice. n=3 mice (*mdx* and *mdx/Pten^cKO^*). (E) Heat map diagram showing the upregulated genes related to ECM and basement membrane in Di muscle of *mdx/Pten^cKO^* versus *mdx* mice. n=3 mice (*mdx* and *mdx/Pten^cKO^*). (F) Western blot analysis of pAkt and sarcolemmal proteins ITGA7, α-SG, α-DG and eMHC in Di muscles of WT, *mdx* and *mdx/Pten^cKO^* mice. (G) Western blot analysis of MG53 in Di muscles of WT, *mdx* and *mdx/Pten^cKO^* mice. (H) Representative immunofluorescence staining of MG53 and α-Laminin on cross-sections of Di muscles from WT, *mdx* and *mdx/Pten^cKO^* mice. Scale bar: 20 μm. (I) Quantification analysis of the relative intensity of MG53 signaling on cross-sections of Di muscles from WT, *mdx* and *mdx/Pten^cKO^* mice. n=4 mice (WT, *mdx* and *mdx/Pten^cKO^*). Data are means ± s.e.m. Two-tailed *t*-test: \**P* < 0.05; n.s., *P* > 0.05.

To unravel the function of differentially expressed genes, we performed Gene Ontology (GO) and Kyoto Encyclopedia of Genes and Genomes (KEGG) pathway analysis using the clusterProfiler software.^31^ The top enriched biological process categories that were upregulated in *mdx/Pten^cKO^* muscle included extracellular matrix, basement membrane, BMP signaling pathway, muscle cell migration, organ growth, muscle tissue development, insulin-like growth factor binding, heart contraction, and wound healing (Figure 5C and Data S3). The enrichments of biological processes related to muscle contraction and membrane integrity further support the observed improvement of muscle function and membrane integrity in *mdx/Pten^cKO^* muscles (refer to Figures 2 and 4). By contrast, the enriched biological processes that were downregulated in *mdx/Pten^cKO^* muscle included small molecular catabolic process, fatty acid metabolic process, lipid biosynthetic process, blood coagulation, acute-phase response, negative regulation of wound healing, complement activation, humoral immune response and acute inflammatory response and so on (Figure 5C and Data S4). The negative enrichments of the coagulation and immune related terms is also consistent with our earlier immunostaining results showing reduced immune cell infiltration and pathology in *mdx/Pten^cKO^* muscle (refer to Figure 3). Similarly, KEGG pathway analysis showed that genes upregulated in *mdx/Pten^cKO^* muscles were significantly related to ECM-receptor interaction and cytokine-cytokine receptor interaction (Figure S5A), whereas genes downregulated in *mdx/Pten^cKO^* muscles were related to retinol metabolism, peroxisome, complement and coagulation cascades, PPAR signaling pathway, and steroid biosynthesis, and so on (Figure S5A).

Interestingly, we found that genes involved in wound healing process were significantly changed in *mdx/Pten^cKO^* muscle (Figure 5C). As a normal biological process, wound healing is achieved through four precisely and highly programmed phases: hemostasis, inflammation, proliferation, and remodeling.^32^ An interference of these processes will cause improper or impaired wound healing. Based on the GO analysis, we generated heat maps to show the expression of genes enriched in biological processes acute-phase response, immune response, ECM and basement membrane, and wound healing categories (Figure 5D, 5E, S5B and S5C). A series of genes (*Crp*, *Orm1*, *Orm2*, *Ugt1a1 Saa1*,and CD163) that are associated with acute-phase response were highly expressed in Di muscles of *mdx* mice, but downregulated significantly in *mdx/Pten^cKO^* muscles, with the Log_2_ Fold Change (Log_2_FC) equal to −6.5, −6.2, −4.9, −4.1, −3.2 and −1.2, respectively (Figure S5C and Data S1). Similarly, genes encoding blood coagulation factors (*F2*, *F11* and *Apoh*) were also downregulated by 5.8, 5.7 and 2.6-Log_2_FC in *mdx/Pten^cKO^* muscles (Figure S5C and Data S1). More than 30 genes related to immune and/or inflammatory response were enriched in *mdx* muscles, but all decreased in *mdx/Pten^cKO^* muscles (Figure S5C and Data S1). Moreover, the genes annotated as negative regulation of wound healing (kininogen, plasminogen and fibrinogen family *Kng1*, *Kng2*, *Plg*, *Fgg*, *Fga* and *Fgb*) were dramatically downregulated in *mdx/Pten^cKO^* muscles with the Log_2_FC equal to −4.1, −2.8, −2.5, −2.5, −2.4 and −1.7, respectively (Figure 5D and Data S1). These unbiased gene expression analyses demonstrate that the Di muscle of *mdx* mice exhibit more severe hemostasis and inflammation phases than *mdx/Pten^cKO^* muscles.

In contrast, several positive regulators of wound healing (*Prkg1*, *Pdgfb* and *Wisp1*) were upregulated significantly in *mdx/Pten^cKO^* compared to *mdx* muscles (Figure 5D and Data S2). Notably, *Frem2* and *Frem1*, two genes encoding integral membrane proteins localized to the basement membrane and required for maintenance of membrane integrity, were significantly upregulated in *mdx/Pten^cKO^* compared to *mdx* muscles, with the Log_2_FC equal to 1.5 and 1.1, respectively (Figure 5E and Data S2). Other upregulated genes in *mdx/Pten^cKO^* muscles including *Col22a1*, *Col15a1*, *Col7a1*, *Itga6*, *Col6a3*, *Lama4*, *Lamb1* and *Adamts17* were all well-known genes encoding basement membrane or ECM-associated components (Figure 5E and Data S2). Proteins encoded by these genes are necessary for maintaining membrane integrity and strengthening skeletal muscle attachments during contractile activity. In addition, the expression of *Ttn*, a protein involved in the assembly and function of muscle sarcomeres, and *Myh2*, functioned in skeletal muscle contraction, were significantly upregulated in *mdx/Pten^cKO^* compared to *mdx* muscles (Data S2). These results suggest that *mdx/Pten^cKO^* muscles exhibit more active ECM formation and remodeling processes compared to the *mdx* muscles. Thus, loss of *Pten* promotes the repair of damaged muscle and restores membrane integrity.

### Upregulation of Membrane Repair Signaling in *mdx/Pten^cKO^* Muscle

As genes involved in the wound healing and membrane integrity pathways are upregulated in *mdx/Pten^cKO^* muscles, we further examined if the protein levels were increased correspondingly. We first examined the protein levels of several sacrolemmal components to confirm the changes of ECM and basement membrane in *mdx/Pten^cKO^* mice. Western blot analysis showed that the phosphorylated Akt (pAkt) was decreased in Di muscle of *mdx* relative to WT mice, but was dramatically elevated in *mdx/Pten^cKO^* mice (Figure 5F). Associated with sarcolemma impairment, the protein levels of integrin α7 (ITGA7), α-sarcoglycan (α-SG), and eMHC were decreased in Di muscle of *mdx* mice (Figure 5F). While *Pten* KO robustly upregulated the expression of ITGA7, α-SG, α-dystroglycan (α-DG), and eMHC in Di muscles of *mdx* mice (Figure 5F). These results validated that *Pten* deletion activated the Akt signaling in *mdx* mice and promoted the ECM remodeling.

Recent studies have identified MG53 as the key components of sarcolemmal membrane repair machinery.^33,34^ MG53 mediates the active trafficking of intracellular vesicles to damaged sites of sarcolemma, and repairs the sarcolemmal lesion and remodels the glycoprotein complex and ECM.^35–37^ As a readout protein indicating membrane repair ability, we next examined the expression of MG53 in *mdx* mice. Interestingly, we found that the protein level of MG53 was downregulated in Di muscle of *mdx* relative to WT mice (Figure 5G), indicating a lower membrane repair capacity in *mdx* muscles. However, the protein level of MG53 was obviously upregulated in *mdx/Pten^cKO^* compared to *mdx* muscles (Figure 5G). Despite the constitutive expression of MG53 protein in WT muscle, MG53 was rarely localized to the trafficking vesicles around myofiber membrane (Figure 5H), potentially due to the lack of damaging stimuli. Intriguingly, MG53 immunofluorescence signal was predominantly localized diffusely inside the myofibers in *mdx* muscle instead of around myofiber membrane (Figure 5H), which again indicating a failure of activating the membrane repair machinery in *mdx* muscle. On the contrary, abundant MG53 signal was observed to co-localize with the trafficking vesicles adjacent to the membrane in *mdx/Pten^cKO^* muscles (Figure 5H). Quantitative analyses showed that the relative intensity of MG53 signal was 2-fold in *mdx/Pten^cKO^* muscles of that in *mdx* muscles (Figure 5I). These results indicate that *Pten* deletion in *mdx* mice might promote MG53 membrane repair machinery.

In support of the above results, we observed an increase in the number of caveolae in *mdx/Pten^cKO^* when compared to *mdx* myofibers (Figure S6A). Caveolae are lipid-enriched microdomains at the plasma membrane that have been shown to participate in numerous cellular processes such as signal transduction, endocytosis, and membrane repair^38^. We found through TEM that the number of caveolae structure per micrometer of plasma membrane was significantly decreased in *mdx* muscle, with 35% fewer than that of WT muscle (Figure S6B). Interestingly, *mdx/Pten^cKO^* doubled the abundance of caveolae in *mdx* muscles (3 versus 1.5 in *mdx/Pten^cKO^* and *mdx* muscle, respectively, Figure S6B), to a level similar to the WT. Together, *Pten* KO restores the integrity of muscle basement membrane accompanying with the upregulation of membrane repair signaling in *mdx/Pten^cKO^* muscle.

### Pharmacological Inhibition of Pten Ameliorates Muscle Pathology and Improves Motor Function of *mdx* Mice

To explore the therapeutic potential of targeting Pten signaling in dystrophic muscles, we treated *mdx* mice with VO-OHpic trihydrate (VO-OHpic), a specific pharmacological inhibitor of Pten that has been validated in several preclinical disease models^39,40^. We intraperitoneally injected littermate *mdx* mice at 3 week-old with VO-OHpic or vehicle control for 21 days consecutively (Figure 6A). We confirmed the Pten inhibition by examining the expression of its downstream Akt pathway. Although pAkt levels varied, the ratio of pAkt to total Akt was consistently increased in the Qu and Gas muscles of VO-OHpic-treated *mdx* mice compared to vehicle-treated *mdx* mice (Figure S7A). We observed no significant changes in body weights and growth curves between VO-OHpic-treated and vehicle-treated *mdx* mice (Figure S7B), indicating negligible side effects by the consecutive treatment of VO-OHpic. Additionally, no significant changes of muscle weight were found between VO-OHpic-treated and vehicle-treated *mdx* mice (Figure S7C).

**Figure 6.**
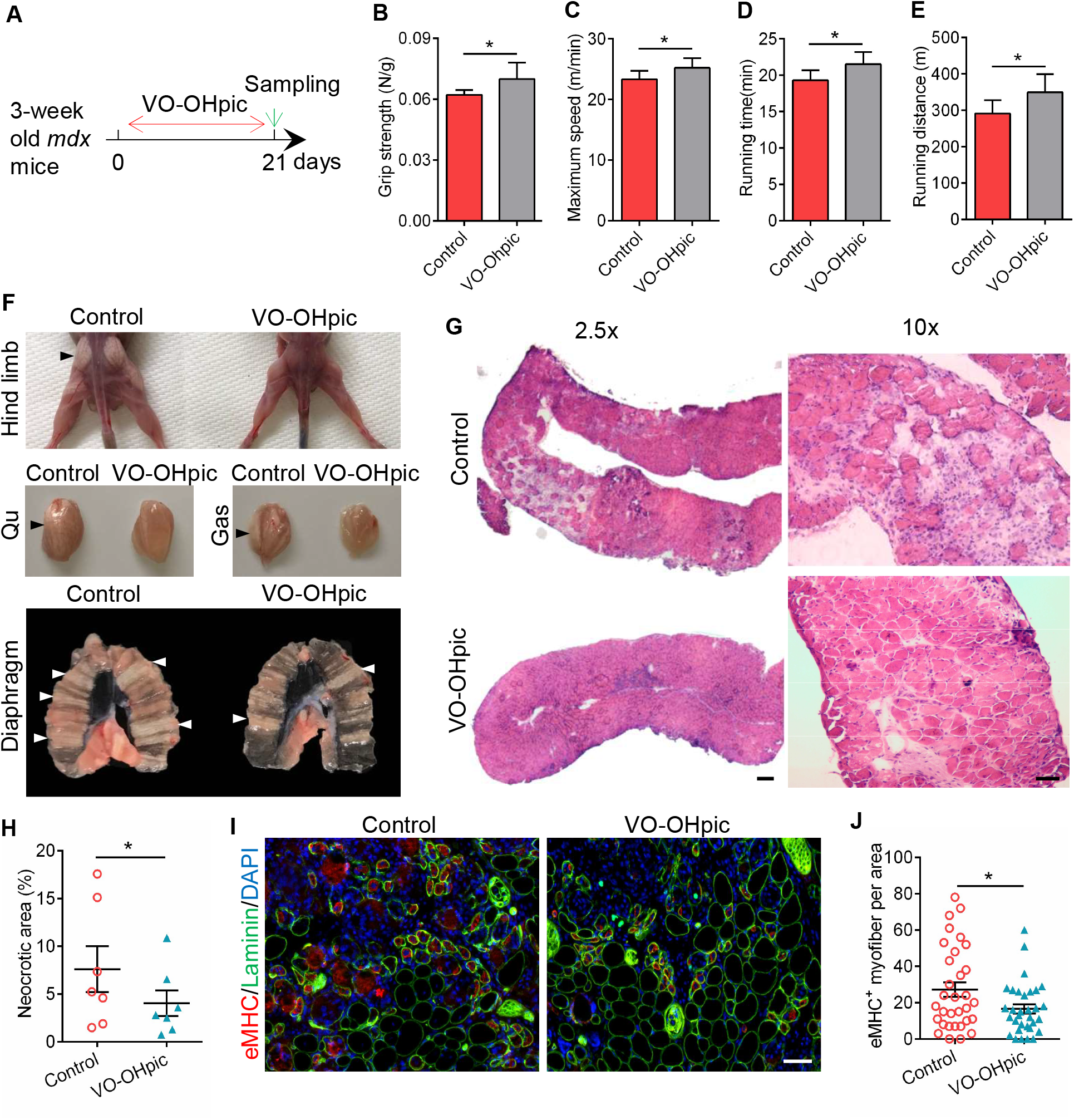
Pharmacological Inhibition of Pten Ameliorates the Symptoms of *mdx* Mice. (A) Schematic showing the treatment procedure of Pten inhibitor VO-OHpic in *mdx* mice at 3 weeks of age. (B) Grip strength test of *mdx* mice 21 days post VO-OHpic treatment. n=11 mice (Control and VO-OHpic). (C-E) Maximum speed (C), total running time (D), and running distance (E) of *mdx* mice 21 days post VO-OHpic treatment. n=12 mice (Control and VO-OHpic). (F) Representative image of hind limb, Qu, Gas, and Di muscles of *mdx* mice mice 21 days post VO-OHpic treatment. n=12 mice (Control and VO-OHpic). (G) Representative HE staining image of Di muscle cross-sections of *mdx* mice at 2.5 and 10 x magnifications 21 days post VO-OHpic treatment. n=12 mice (Control and VO-OHpic). Scale bar: 100 μm for 2.5x, 50 μm for 10x. (H) Quantification analysis of necrotic area in Di muscles of *mdx* mice 21 days post VO-OHpic treatment. n=7 mice (Control and VO-OHpic). (I) Representative immunofluorescence staining of eMHC and α-Laminin on Qu muscle cross-sections of *mdx* mice 21 days post VO-OHpic treatment. Scale bar: 50 μm. (J) Quantification analysis of the number of eMHC^+^ cells on Qu muscle cross-sections of *mdx* mice 21 days post VO-OHpic treatment. n=4 mice (Control and VO-OHpic), each dot represents one image and total 32 images were took from 4 mice for each group. Data are means ± s.e.m. Two-tailed *t*-test: \**P* < 0.05.

We then investigated the muscle function of the *mdx* mice by employing grip strength and treadmill test. Notably, VO-OHpic-treated *mdx* mice displayed better grip strength (Figure 6B), maximum running speed (Figure 6C), total running time (Figure 6D) and running distance (Figure 6E) compared to vehicle-treated *mdx* mice, suggesting significant improvements of muscle function after the Pten inhibitor treatment. Further morphological analysis showed that the fibrotic tissues were reduced considerably in the whole hind limb and Di muscles of *mdx* mice after VO-OHpic treatment, compared to the severe fibrosis in vehicle-treated *mdx* mice (Figure 6F). Histologically, remarkable reduction of the necrotic areas was observed in Di, Gas, and Qu muscles of VO-OHpic-treated *mdx* mice (Figures 6G and S7D). Strikingly, VO-OHpic-treated *mdx* mice exhibited 47 % and 61 % decreases in the necrotic areas in Di, and Qu muscles, respectively, compared to that of vehicle-treated *mdx* mice (Figures 6H and S7E). In addition, we found that the numbers of eMHC^+^ cells were reduced significantly in Gas and Qu muscles of VO-OHpic-treated *mdx* mice (Figures 6I, 6J, S7F and S7G), which suggests the diminished degenerating and regenerating activities in dystrophic muscles after Pten inhibitor treatment. Taken together, these results reveal that the pharmacological inhibition of Pten improves the dystrophic muscle function and attenuates pathophysiology of *mdx* mice.

## Discussion

Our current study reveals a previously unrecognized critical role of Pten signaling in regulating the dystrophic muscle function and pathogenesis of the *mdx* mouse. We provided compelling genetic, physiological, histological, pathological and molecular evidence to demonstrate that *Pten* deletion robustly restores muscle function and diminishes the symptoms in *mdx* mice. Loss of *Pten* prevents dystrophic myofiber degeneration by maintaining sarcolemmal integrity and promotes the repair of damaged myofiber membrane, accompanied with the activation of Akt and MG53 signaling, which eventually benefits the dystrophic muscle function. Importantly, pharmacological inhibition of Pten mimics the phenotypes of *Pten* deletion in *mdx* mice; both significantly improve muscle function and ameliorate the DMD symptoms. Overall, our study provides promising evidence that targeting Pten represents a novel therapeutic strategy for DMD treatment.

The primary muscle pathology caused by dystrophin mutation includes continuous myofiber degeneration, muscle wasting and weakness due to the disruptions of the lineage among extracellular matrix, myofiber membrane and intracellular actin cytoskeleton.^29,41–43^ The elevated Pten level was previously reported in DMD patients and animal models of severe muscle wasting and weakness.^18–21^ Our observations reveal for the first time the critical role of Pten signaling in DMD pathogenesis. Impressively, Pten inhibition significantly diminishes the progressive muscle wasting and weakness in *mdx* mice, which is consistent with previous studies that constitutive activation Akt or inactivation of Pten regulators ameliorates pathology of dystrophic muscle and improves function.^18,22,23,44,45^ Importantly, our observations show increased myofiber membrane integrity in line with ECM proteins expression, and upregulated wound healing pathways in *mdx* mice upon *Pten* deletion. Given loss of *Pten* promotes muscle growth and results in muscle hypertrophy in normal mice,^24^ our findings suggest that Pten inhibition might benefits the dystrophic muscles through two potential mechanisms as that resisting the dystrophin deficiency-induced muscle damage and repairing the myofibers upon damages. These findings provide a strong rationale for the identification of mechanisms to boost muscle regrowth to counter the muscle pathologies in muscular dystrophy.

Cell membrane integrity and repair is a fundamental aspect of normal physiology and is critical for maintaining cellular homeostasis.^36–38^ The disruption of this integrity and repair function underlies the progression of many human diseases such as muscular dystrophy.^1^ Emerging evidence points to the possibility that restoration of dystrophic muscle function by strengthening the mechanical stability of sarcolemma, or by modulating sarcolemmal membrane repair machinery.^15,46–50^ MG53 and its co-regulators dysferlin, Annexin and Cav3 have been identified as the key components of sarcolemmal membrane repair machinery in skeletal muscle.^33,34,51,52^ Ablation of either of them in mice results in severe muscular dystrophy,^34,52,53^ while overexpression of them boosts sarcolemmal membrane repair and protects against damage in a mouse model of muscular dystrophy.^54–57^ Although *Pten* KO did not alter the mRNA levels of MG53, it upregulated the protein levels of MG53 in the muscle. Importantly, in parallel to the increase of caveolae numbers, MG53 translocate to vesicles in *Pten* KO muscles, rather than the diffuse location observed in *mdx* myofibers. This indicates that Pten signaling might regulate MG53 synthesis or translocation to trafficking vesicles translationally or post-translationally. Certainly, one essential study in future is to elucidate how MG53 translocation is blocked in *mdx* muscle and how Pten signaling mediates the translocation of MG53 to vesicles that trafficking to the damaged sites.

We used the new D2-*mdx* mouse strain to better model DMD pathology. The traditional C57BL/10*-Dmd^mdx^/J* (B10-*mdx*) mouse model has been the most commonly used preclinical animal model to study DMD.^26–28^ However, a fatal caveat of the B10-*mdx* mouse model is that its pathological features are much less severe compared to DMD patients, manifested by the very late onset of myopathology, increased muscle mass, hypertrophic myofibers, the absence of fibrosis, and the normal regenerative capacity due to enhanced self-renew activity of muscle stem cells. These features strongly limit the translational value of the B10-*mdx* mouse model.^26–28^ The D2-*mdx* mouse strain better recapitulates the human characteristics of DMD myopathology including earlier onset of pathology, lower hind limb muscle weight, increased fibrosis and remarkable muscle weakness.^26–28^ We predict that the current findings with this D2-*mdx* mouse model will be readily translated to treat DMD in humans.

Our findings suggest that targeting Pten signaling represents a potential therapeutic strategy that could be translated into the clinical treatment of DMD patients. Indeed, the overall positive role of the PI3K/AKT signaling pathway in cell growth and survival underlies the rational for the therapeutic targeting of PTEN in tissue regenerative medicine.^39,40,58^ The potential side effect of targeting Pten to treat DMD is the risk of tumorigenesis in other non-muscle tissues,^39,40,58^ although no increased incidence of spontaneous tumors was observed in *mdx* mice with skeletal muscle *Pten* deletion within the present experimental period. Nevertheless, methodologies that specifically target Pten in dystrophic muscles are needed to avoid the side effect of Pten inhibition with aging. The advances in nanotechnology and controlled drug delivery enable further development of the therapeutic strategy.^30,59^ For example, ligand-targeted nanoparticles have displayed unique advantages for effective and tissue-specific drug delivery, and have shown promising therapeutic efficacy for the treatment of cancer,^60^ neurodegenerative disease,^61^ and obesity.^62^ Moreover, recent efforts on identification of muscle targeting peptides offered the potential for specific drug delivery to DMD muscles by peptide-functionalized nanoparticles.^63,64^ Therefore, future efforts in development of a nanotechnology-based drug delivery system to targeting Pten in dystrophic muscles is warranted in human clinical trials.

## Materials and Methods

### Mice

The mouse strains *MyoD^Cre^* (#014140), *Pten^f/f^* (#006440), and *Dmd^mdx^* (#013141) used in this study were purchased from Jackson Laboratory. Genotyping of the mouse was performed by genomic DNA isolated from ear, and were screened by PCR using primers and program following the protocols provided by the company. The genotypes for *Pten* knockout and control *mdx* mice as well as wild type under D2.B10 background are as follows: *mdx* (♀: *Pten^f/f^*;*Dmd^mdx^/Dmd^mdx^*, ♂: *Pten^f/f^;Dmd^mdx^/Y*), *mdx/Pten^cKO^* (♀: *MyoD^Cre^*;*Pten^f/f^*;*Dmd^md^/Dmd^mdx^*, ♂: *MyoD^Cre^*;*Pten^f/f^;Dmd^mdx^/Y*), and wild-type (*Pten^f/f^*). For some experiments, wild-type mice under C57BL/6J background were used as control. The mice were housed in a 12-h light–dark cycle and given water and food ad libitum. All animal experimental protocols were approved by the Purdue Animal Care and Use Committee. Generally 7- to 9-week-old mice were used for all experiments as stated specifically in figure legends. Both male and female mice were used for all studies and always gender-matched for each specific experiment.

### Pten Inhibitor Treatment

For pharmacological inhibition of Pten, Pten inhibitor VO-OHpic trihydrate (Sigma-Aldrich) was dissolved in 2% dimethyl sulfoxide water solution containing 40% PEG300 and 2% Tween 80 with a final concentration of 1 mg/mL. VO-OHpic trihydrate was intraperitoneally injected into 3 week-old *mdx* mice at the dosage of 10 mg/kg body weight for 21 days consecutively, with the littermate *mdx* mice injected with vehicle (2% dimethyl sulfoxide water solution containing 40% PEG300 and 2% Tween 80) as control. Mice were subjected to grip strength and treadmill test, and sacrificed at the same day of the test.

### Whole Body Composition Analysis

The fat and lean mass compositions of mice were analyzed with Echo-MRI 130 analyzer (EchoMRI LLC, Houston, TX, USA). Briefly, the instrument was first calibrated with corn oil. Parameters including total body fat, lean mass, free water, and total body water, were measured in a rapid (approximately 30 seconds) and noninvasive manner with three replicates for each measurement. The measurement tubes were disinfected with 70% ethanol between testing each mouse.

### Treadmill Exercise Performance Test

The exercise tolerance test was performed using an Exer-3/6 treadmill (Columbus Instruments). Mice were first trained for 3 consecutive days before the test to achieve adaptation (10 min per day at 10 m/min, +10° slope and 3 Hz+1 mA electrical stimulation). For the test, mouse was placed in a dedicated lane and started to run on the treadmill at 10 m/min for 5 min, with the gradual increased speed by 2 m/min every 2 min until they were exhausted. Exhaustion was determined when the mouse was unable to run on the treadmill for 10 s upon 5 times pushing. The maximum speed, total running distance and running time achieved were recorded by Treadmill software (Columbus Instruments).

### Whole-limb Grip Strength Test

A digital grip-strength meter (Columbus Instruments) was used to measure four-limb grip strength in mice following the manufacture’s guide. Briefly, mice were acclimatized for 10 min before the test in the hood. The mouse was allowed to grab the metal pull bar with the paws. Then the mouse tail was gently pulled backward until mice could not hold on the bar. The force at the time of release was recorded as the peak tension. Each mouse was tested ten times. The grip strength was defined as the average strength measured with or without normalization with bodyweight.

### Assessment of Muscle Contractile Function

Muscle contractile force measurement was conducted by using an *in vitro* muscle test system (1200A Intact Muscle Test System, Aurora Scientific).^65^ In brief, the mice were anesthetized with isoflurane, the hind limb was excised and immediately placed in a bicarbonate-buffered solution (137 mM NaCl, 5 mM KCl, 1 mM MgSO_4_, 1 mM NaH_2_PO_4_, 24 mM NaHCO_3_, and 2 mM CaCl_2_) equilibrated with 95% O_2_-5% CO_2_ (pH7.4) for dissection. The EDL muscle was dissected first and subjected to the assessment of contractile function, followed by the soleus muscle. After isolation, the proximal and distal tendons were tied with braided silk suture thread (4-0, Fine Science Tools). The muscle was then mounted in the muscle test system continuously bubbled with carbogen (5% CO_2_ in O_2_) at room temperature. The stimulation protocol consisted of supramaximal electrical current delivered through platinum electrodes using a biphasic high-power stimulator (701C; Aurora Scientific). After determining optimum length (Lo) with a series of twitch stimulations at supramaximal voltage (optimal length, *L*o), the temperature of the organ bath was increased to 32°C. After 10 min of thermoequilibration, a force-frequency relationship was generated using select frequencies between 1 and 300 Hz for the EDL muscle (200-ms train) and 1 and 200 Hz for the soleus muscle (500-ms train). This was followed by a 5-min fatiguing protocol of 500-ms volleys of 40-Hz stimulation applied once every 2 s. The muscle was then removed from the organ bath, trimmed of connective tissue, blotted dry, and weighted. Muscle cross-sectional area (CSA) was determined by dividing the wet muscle mass by the product of *L*o and muscle-specific density (1.056 g/cm^3^). Specific force (N/cm^2^) was calculated by dividing the muscle force (N) by the CSA (cm^2^). The maximum twitch response was analyzed for peak twitch tension, time to peak twitch tension, and twitch half-relaxation time.

### Single Myofiber Isolation

Single myofibers were isolated from extensor digitorum longus (EDL) muscles by 1 hour digestion in 0.2% collagenase type I (Worthington Biochemical Corporation) in Dulbecco’s Modified Eagle’s Medium (DMEM) at 37 °C with gently rotation. The digested muscle was then transferred to horse serum-coated culture dishes (60-mm) containing pre-warmed DMEM with 1% penicillin–streptomycin. Single myofibers were isolated by gently flushing muscles with a fire-polished glass pipette. Individual myofibers were then collected by pipette to remove dead myofibers and cellular debris, and subjected to protein extraction and western blot analysis.

### Creatine Kinase (CK) Activity Test

Blood was collected from the heart of mice immediately following sacrifice and left at room temperature for 30 min to achieve coagulation. Serum was then separated from other blood fractions by centrifugation at 1,000 g for 10 min and stored at −80 °C for further use. CK activity in the serum was measured using Creatine Kinase Activity Assay Kit (MAK116, Sigma).

### Evans Blue Uptake

Evans blue (EB) uptake analysis was conducted to check the muscle membrane integrity of WT, *mdx* and *mdx/Pten^cKO^* mice. In detail, EB (10 mg/mL in PBS) was administered to mice intraperitoneally (0.04 mL per 10 g body weight). After 24 h, Di, Gas and Qu muscles were harvested and sectioned using a cryostat (Leica CM 1850). EB was detected as red fluorescence under the fluorescent microscope.

### Histology and Immunofluorescence Staining

Whole muscle tissues (TA, Gas, Qu, and DI) from WT, *mdx* and *mdx/Pten^cKO^* mice were dissected and frozen immediately in an O.C.T. (optimum cutting temperature) compound. Frozen muscles were sectioned using Leica CM1850 cryostat at 10 μm thickness. Slides were subjected to histological staining or immunofluorescence staining. For hematoxylin and eosin (H&E) staining, the slides were stained in hematoxylin solution for 20-30 min, rinsed in running water for 3 times. Slides were then stained in Eosin solution for 1-2 min, dehydrated in graded ethanol and Xylene, and covered using Permount for further imaging. For Masson’s trichrome staining, slides were fixed for 15 minutes at room temperature in a formalin/ethanol mixture (40 mL of 100% ethanol to 10 mL of 37% formaldehyde), then rinsed in water and trichrome stain carried out. Slides were stained in weigert’s hematoxylin 10 minutes, rinsed in tap water for bluing, stained biebrich scarlet-acid fuchsin solution 3-5 minutes, differentiated in phosphotungstic-phosphomolybdic acid until collagen was decolorized and counterstained in aniline blue solution for less than 1 minute. Slides were dehydrated, cleared and cover-slipped in a Leica Autostainer XL.

For immunofluorescence staining, cross-sections were fixed in 4% paraformaldehyde (PFA) in PBS for 10 min and incubated in blocking buffer (5% goat serum, 2% BSA, 0.1% Triton X-100, and 0.1% sodium azide in PBS) for 1 hour at room temperature. Samples were then incubated with primary antibodies anti-α-Laminin (L9393, Sigma, 1:1,000), anti-α-Laminin (ab2466, abcam, 1:1,000), anti-eMHC (F1.652, DSHB, 1:100), Alexa Fluor 647 anti-mouse CD68 (FA-11, BioLegend, 1:1,000), or anti-mouse MG53 (gift from Jianjie Ma, The Ohio State University, 1:500) at 4 ℃ overnight. After incubated with corresponding second antibodies and DAPI at room temperature for 1 hour, samples were cover with fluorescent mounting medium for further imaging.

All histological images were captured using a Nikon D90 digital camera mounted on a microscope with ×20 objectives. All immunofluorescent images were captured using Leica DM 6000B microscope with ×20 objectives. Images for WT, *mdx* and *mdx/Pten^cKO^* mice samples were captured using identical parameters. All images shown are representative results of at least three biological replicates. The cross-sectional area (CSA) and feret diameter of myofiber were measured based on α-Laminin staining. Necrotic area was indicated by DAPI staining defined as the area of mononuclear aggregating. Fibrotic area was defined by Masson’s trichrome staining. Quantifications for all of the histological and immunofluorescence was performed by ImageJ software.

### Transmission Electron Microscopy (TEM) and Membrane Integrity Quantification

Briefly, isolated muscle tissue (cut into 1mm blocks if the muscle was large) was fixed in fixative buffer (2.5% glutaraldehyde, 1.5% paraformaldehyde in 0.1M cacodylate buffer). Samples were rinsed in deionized water followed by fixation in 2% osmium tetroxide for 1 h. Then, the samples were washed in deionized wat water followed by fixation in 1% uranyl acetate for 15 minutes. After washing with deionized wat water, the samples were dehydrated with series of graded ethanol followed by dehydration in acetonitrile. The samples were embedded in epoxy resin (EMbed 812: DDSA: NMA 5:4:2; 0.22 DMP-30). Ultrathin section (longitudinal section) were cut at 70 nm and stained with uranyl acetate and lead citrate. Stained sections were examined under Tecnai T12 transmission electron microscope attached with a Gatan imaging system under 6,000× and 26,000× magnifications. Membrane integrity of myofibers was quantified based on the TEM images. High integrity membrane of myofiber was defined when both the extracellular matrix and sarcolemma layer were observed. In contrast, low integrity membrane of myofiber was defined when the extracellular matrix or sarcolemma layer was missed in the myofiber.

### Protein Extraction and Western Blot Analysis

Total protein was isolated from muscle tissues with RIPA buffer containing 25 mM Tris-HCl (pH 8.0), 150 mM NaCl, 1mM EDTA, 0.5% NP-40, 0.5% sodium deoxycholate and 0.1% SDS. Protein concentrations of samples were determined using Pierce BCA Protein Assay Reagent (Pierce Biotechnology). Total protein extract was resolved by SDS-PAGE, and transferred onto polyvinylidene fluoride membrane (Millipore Corporation). The membranes were incubated in blocking buffer (5% fat-free milk) for 1 h at room temperature, and then blotted with primary antibodies anti-phospho-Ser473 Akt (#4060, Cell Signaling, 1:2,000), anti-total Akt (#4691, Cell Signaling, 1:2,000), anti-Integrin alpha-7 (9.1 ITGA7, DSHB, 1:100), anti-alpha-sarcoglycan (IVD3(1)A9, DSHB, 1:100), anti-alpha-Dystroglycan (IIH6 C4, DSHB, 1:100), anti-eMHC (F1.652, DSHB, 1:100), anti-MG53 (gift from Jianjie Ma, The Ohio State University, 1:1000), and anti-GAPDH (sc-32233, Santa Cruz Biotechnology, 1:5,000) at at 4 ℃ overnight. After incubated with corresponding second antibodies at room temperature for 1 hour, membranes were incubated with chemiluminescence substrate (Santa Cruz Biotechnology) and detected using FluorChem R system (Proteinsimple).

### Total RNA Extraction and RNA-seq Analysis

Total RNA was extracted from diaphragm muscles of *mdx* and *mdx/Pten^cKO^* mice using TRIzol reagent according to the manufacturer’s instructions, and subjected to RNA-seq analysis performed by Novel Bioinformatics Co., Ltd. (https://en.novogene.com/). Briefly, RNA quality analysis was checked by Agarose Gel Electrophoresis and Agilent 2100. A complementary DNA library was then constructed using polyA selected RNA, and sequencing was performed according to the Illumina HiSeq standard protocol. Raw reads from RNA-seq libraries are filtered to remove reads containing adapters or reads of low quality. After filtering, statistics analysis of data production and quality was performed to confirm the sequencing quality. Reference genome and gene annotation files were downloaded from a genome website browser (NCBI/UCSC/Ensembl). TopHat2 was used for mapping the filtered reads to the reference genome. For the quantification of gene expression level, HTSeq V0.6.1 was used to analyze the read numbers mapped for each gene. The FPKM of each gene was calculated based on the gene read counts mapped to genes or exons. A differential expression analysis was performed using the DESeq2 R/EdgeR R package with the threshold of significance set as adjusted *p* < 0.05. Volcano plots are used to infer the overall distribution of differentially expressed genes. Cluster analysis was performed with different samples to check the correlation pattern using hierarchical clustering distance method. GO and KEGG pathway enrichment analysis of differentially expressed genes was performed by using the clusterProfiler software.^31^

### Statistical Analysis

The researchers involved in the *in vivo* studies including grip strength, treadmill, muscle contractile force measurements were blinded to group identity. All images were randomly captured from samples and analyzed in a blinded manner. All statistical analyses and graphing were performed using GraphPad Prism 6.0 (GraphPad Software). All experimental data are represented as mean ± s.e.m. Statistical significance was determined by student’s *t*-test under two-tailed with *P* < 0.05 was considered as significant.

## Acknowledgments

We thank Drs. Hua Zhu and Jianjie Ma (The Ohio State University, USA) for providing the MG53 antibody, Zhongyin Zhang (Purdue University, USA) for advice on PTEN inhibitor, Jun Wu for mouse colony maintenance and members of the Kuang laboratory for their kind assistance. We are grateful to Purdue Electron Microscopy Facility for assistance on transmission electron microscopy, and Purdue Histology Research Laboratory for assistance on Masson’s trichrome staining of cryo-section. This work was supported by grants from the Muscular Dystrophy Association to F.Y. (MDA516161), the National Institutes of Health to S.K. (NIH 1R01AR071649) and to M.D. (NIH R03AR068108).

## Supplemental information

Supplemental information includes seven Supplementary Figures and four Supplementary Data.

## Author contributions

F.Y. and S.K developed the concept for the studies. F.Y., C.S. and S.K designed the experiments. F.Y., C.S., D.H. N.N., J.Q., Z.J., Z.Y., S.O. and B.R. performed the experiments and analyzed the data. M.D. provided reagents. F.Y., C.S. and S.K wrote the manuscript.

---

**Figure S1.**
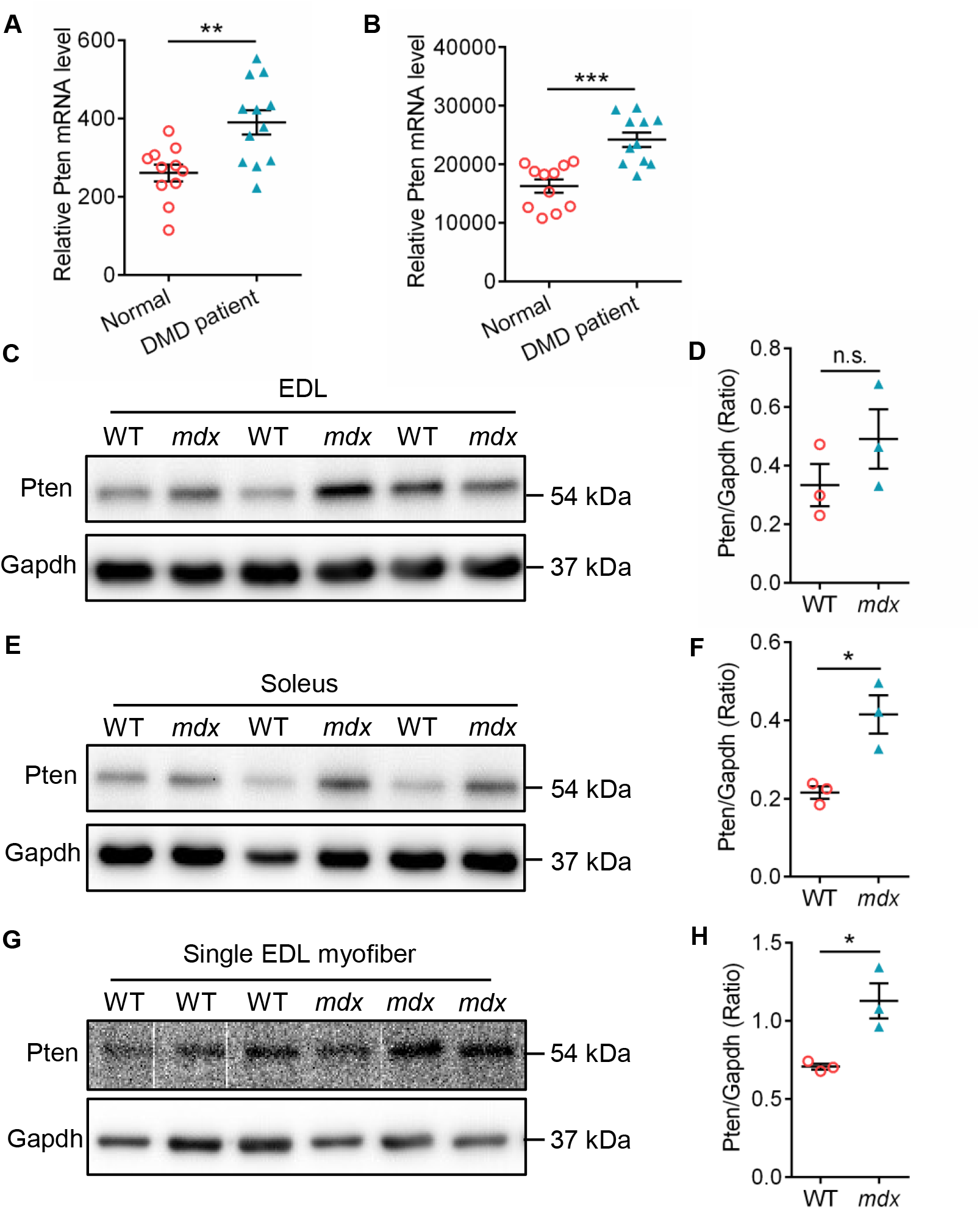
Elevated *PTEN* Expression Level in Skeletal Muscles of Human *DMD* Patient and *mdx* Mice. (A and B) mRNA level of PTEN in quadriceps skeletal muscle biopsies of DMD patient from two online datasets: GSE1004 (A) and GSE1007 (B). (C) Western blot analysis of Pten protein level in EDL muscles of WT and *mdx* mice. (D) Quantification analysis of the ratio of Pten protein level to Gapdh in EDL muscles of WT and *mdx* mice. (E) Western blot analysis of Pten protein level in soleus muscles of WT and *mdx* mice. (F) Quantification analysis of the ratio of Pten protein level to Gapdh in soleus muscles of WT and *mdx* mice. (G) Western blot analysis of Pten protein level in isolated EDL myofiber of WT and *mdx* mice. (H) Quantification analysis of the ratio of Pten protein level to Gapdh in isolated EDL myofiber of WT and *mdx* mice. Data are means ± s.e.m. Two-tailed *t*-test: \**P* < 0.05; \*\**P* < 0.01; \*\*\**P* < 0.001; n.s., *P* > 0.05.

**Figure S2.**
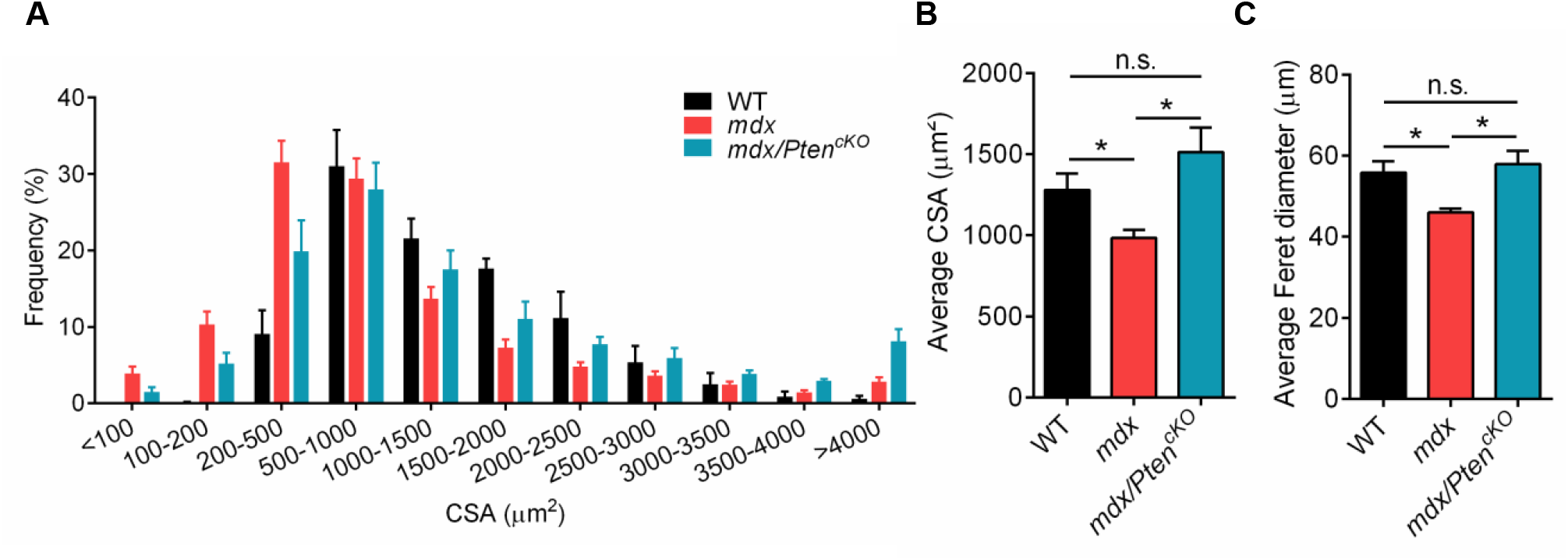
Loss of *Pten* Enlarges Myofiber Size in *mdx* Mice. (A) The frequency of cross-sectional area (CSA) distribution for Gas muscle in WT, *mdx* and *mdx/Pten^cKO^* mice at 9 weeks of age. n=4 mice (WT, *mdx* and *mdx/Pten^cKO^*). (B) The average CSA of Gas muscles in WT, *mdx* and *mdx/Pten^cKO^* mice. n=4 mice (WT, *mdx* and *mdx/Pten^cKO^*). (C) The average Feret diameters of Gas myofber in WT, *mdx* and *mdx/Pten^cKO^* mice. n=4 mice (WT, *mdx* and *mdx/Pten^cKO^*). Data are means ± s.e.m. Two-tailed *t*-test: \**P* < 0.05; \*\**P* < 0.01; \*\*\**P* < 0.001; n.s., *P* > 0.05.

**Figure S3.**
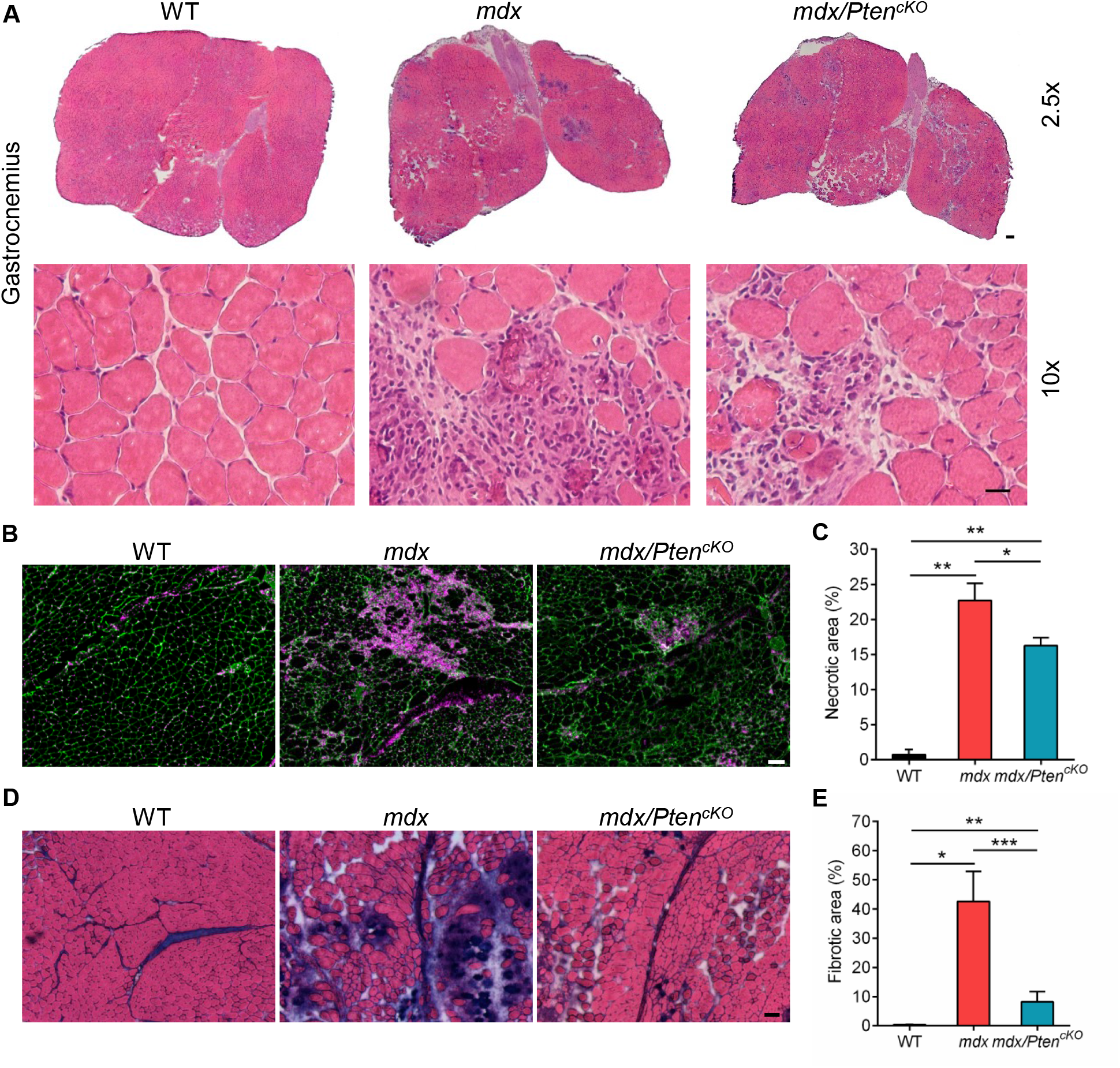
Loss of *Pten* in *mdx* Mice Reduces Necrotic and Fibrotic Myofibers. (A) Representative HE staining image of Gas muscles of WT, *mdx* and *mdx/Pten^cKO^* mice at 9 weeks of age. Scale bar: 2.5x, 200 μm; 10x, 50 μm. (B) Representative images of α-Laminin and DAPI staining showing the necrotic area on cross-section of Gas muscle. Scale bar: 100 μm. (C) Quantification analysis of necrotic area in Gas muscles of WT, *mdx* and *mdx/Pten^cKO^* mice. n=3 mice (WT, *mdx* and *mdx/Pten^cKO^*). (D) Representative images of Masson’s trichrome staining on cross-section of Gas muscle from WT, *mdx* and *mdx/Pten^cKO^* mice. Scale bar: 100 μm. (E) Quantification analysis of the percentage of fibrotic area in Gas muscle from WT, *mdx* and *mdx/Pten^cKO^* mice. n=3 mice (WT, *mdx* and *mdx/Pten^cKO^*). Data are means ± s.e.m. Two-tailed *t*-test: \**P* < 0.05; \*\**P* < 0.01; \*\*\**P* < 0.001.

**Figure S4.**
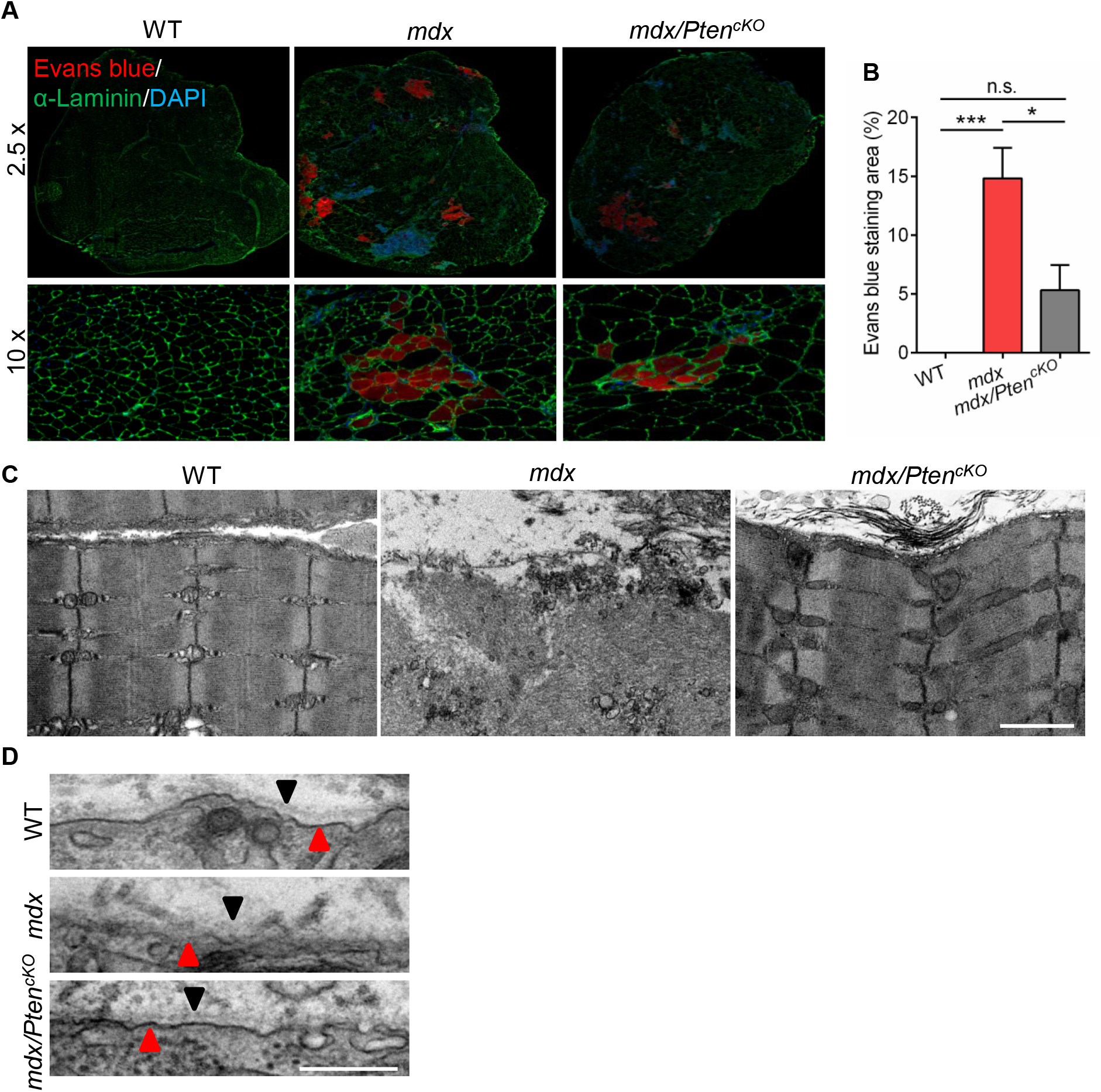
Increased Membrane Integrity in Skeletal Muscles of *mdx/Pten^cKO^* Mice. (A) Representative Evans blue staining images of Qu muscle cross-section from WT, *mdx* and *mdx/Pten^cKO^* mice at 9 weeks of age. (B) Quantification analysis of Evans blue staining area in cross-section of hind limb muscles. n=3 mice (WT, *mdx* and *mdx/Pten^cKO^*). (C) Representative TEM images of EDL muscle longitudinal-sections from WT, *mdx* and *mdx/Pten^cKO^* mice at 8 weeks of age. Scale bar: 500 nm. (D) Representative TEM images of Qu muscle longitudinal-sections from WT, *mdx* and *mdx/Pten^cKO^* mice. Scale bar: 250 nm. Data are means ± s.e.m. Two-tailed *t*-test: \**P* < 0.05; \*\*\**P* < 0.001; n.s., *P* > 0.05.

**Figure S5.**
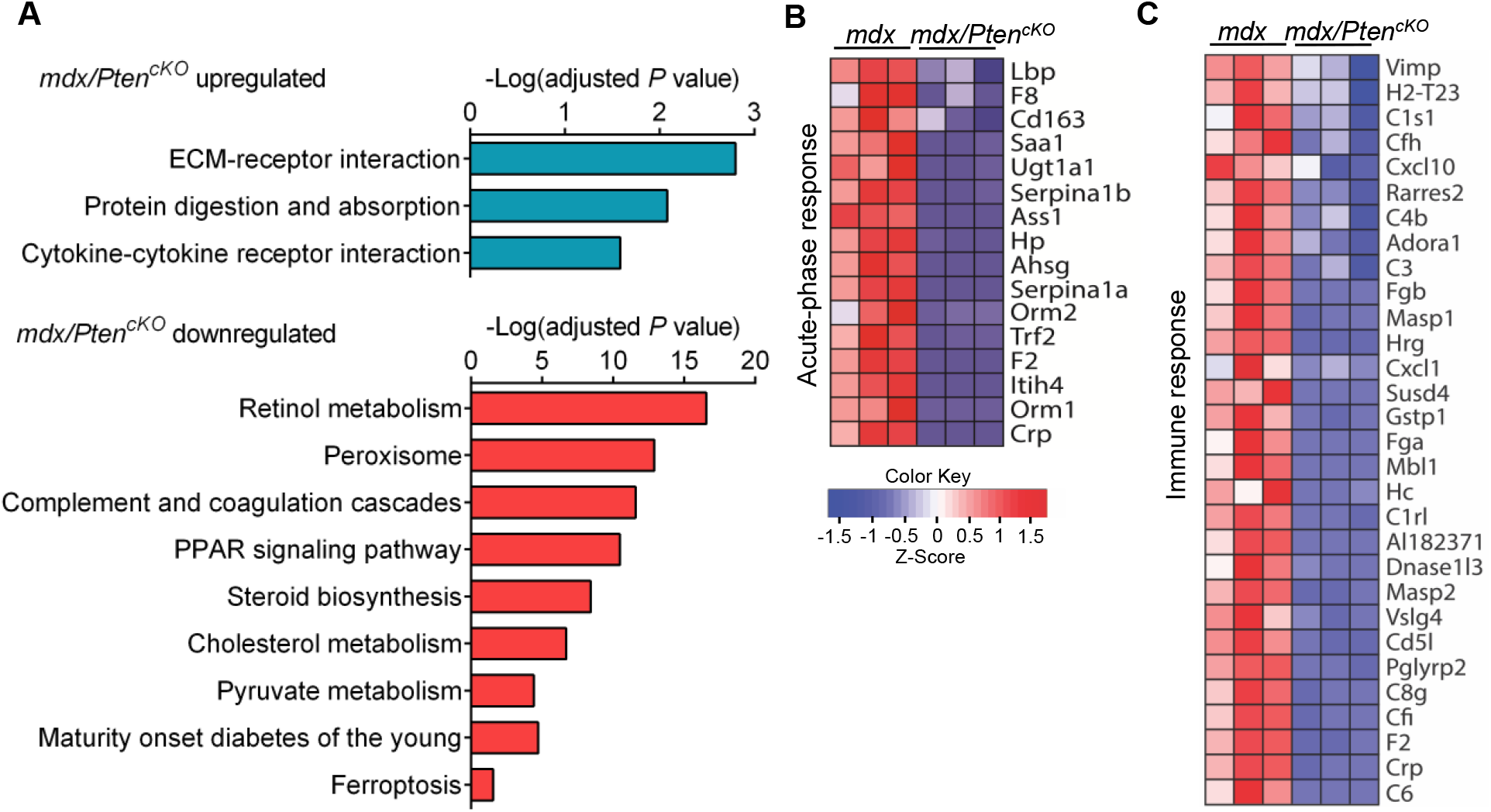
KEGG Pathway Enrichments and Heat Maps in *mdx/Pten^cKO^* Muscle. (A) KEGG analyses showing the enrichment of pathways in Di muscle of *mdx/Pten^cKO^* versus *mdx* mice. n=3 mice (*mdx* and *mdx/Pten^cKO^*). (B) Heat map diagram showing the downregulated genes related to acute phase response in Di muscle of *mdx/Pten^cKO^* versus *mdx* mice. n=3 mice (*mdx* and *mdx/Pten^cKO^*). (D) Heat map diagram showing the downregulated genes related to immune response in Di muscle of *mdx/Pten^cKO^* versus *mdx* mice. n=3 mice (*mdx* and *mdx/Pten^cKO^*).

**Figure S6.**
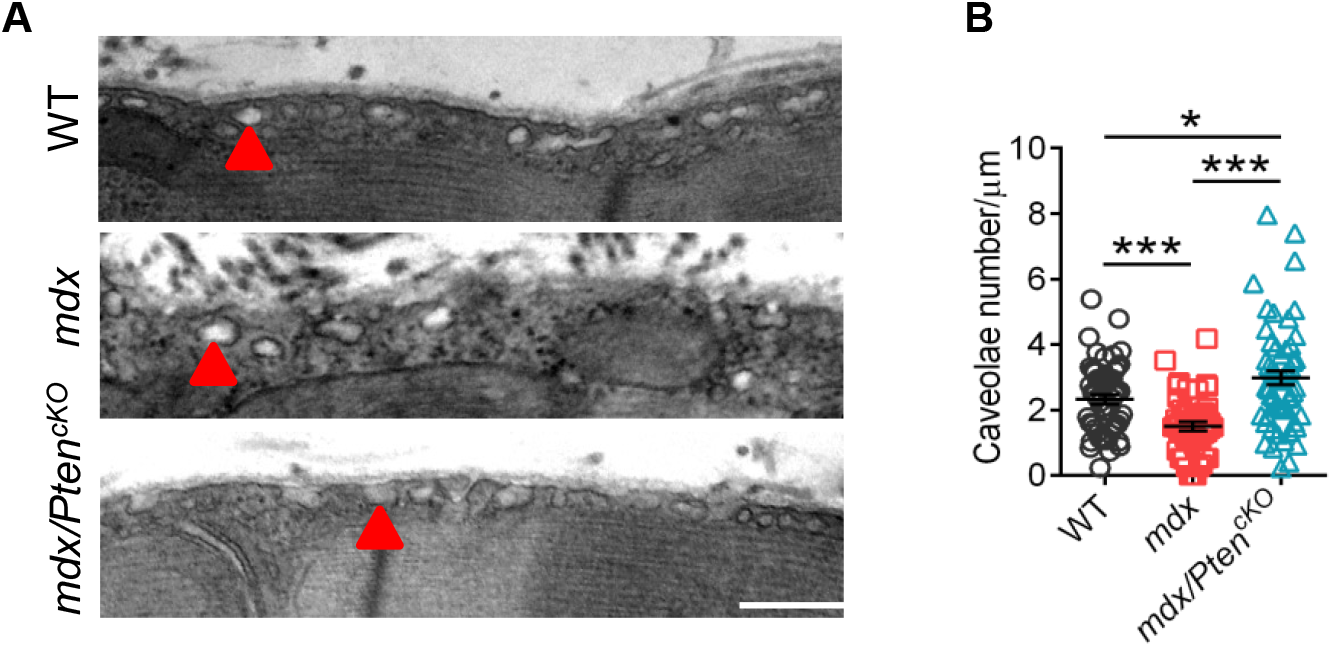
Increased Caveolae Number in Skeletal Muscles of *mdx/Pten^cKO^* Mice. (A) Representative TEM images of EDL muscle longitudinal-sections from WT, *mdx* and *mdx/Pten^cKO^* mice. Scale bar: 500 nm. (B) Quantification analysis of the caveolae number based on TEM images of EDL muscle from WT, *mdx* and *mdx/Pten^cKO^* mice. n=3 mice (WT, *mdx* and *mdx/Pten^cKO^*), Each dot represents one image. Data are means ± s.e.m. Two-tailed *t*-test: \**P* < 0.05; \*\*\**P* < 0.001; n.s., *P* > 0.05.

**Figure S7.**
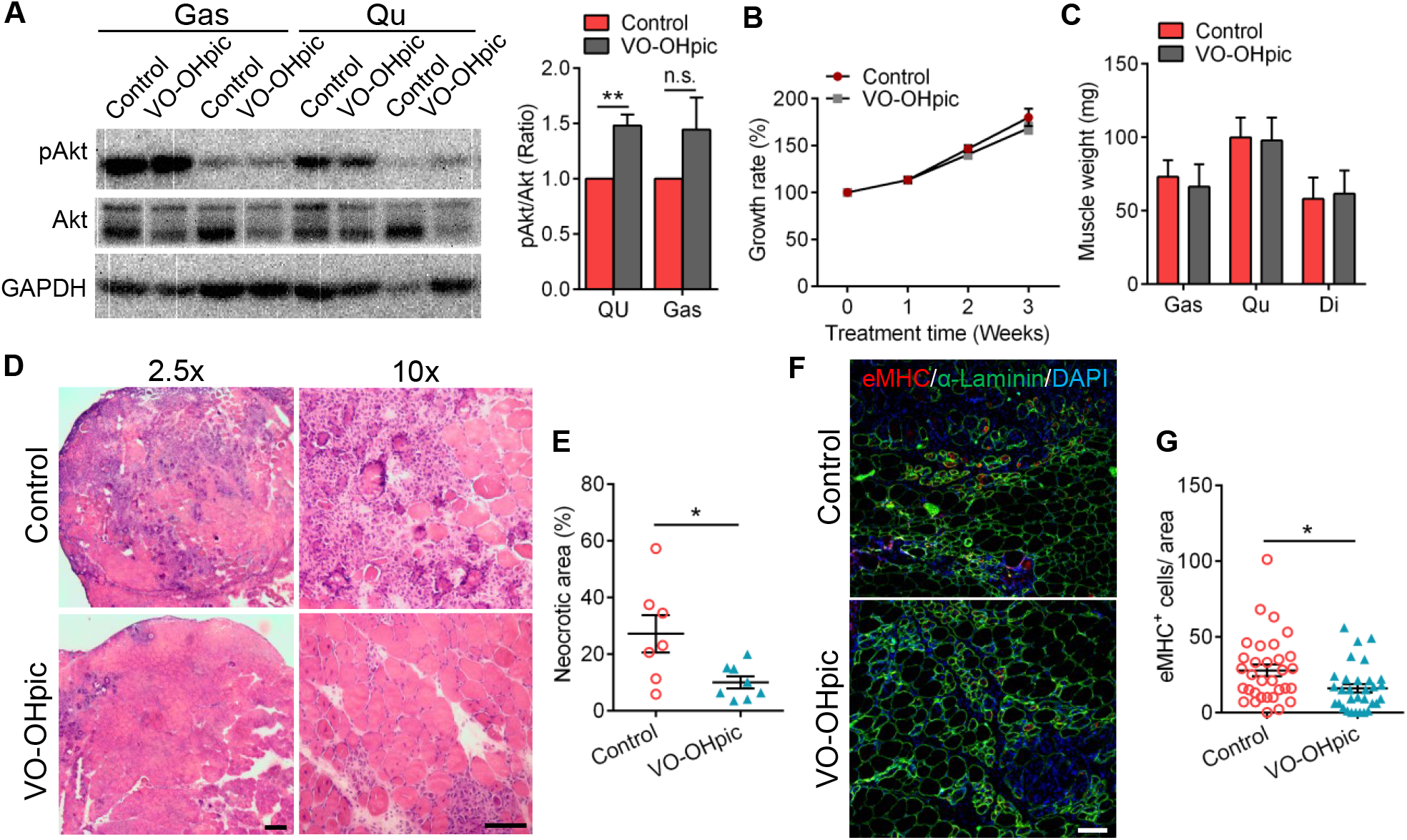
Treatment of Pten Inhibitor Ameliorates Dystrophic Pathology of *mdx* Mice. (A) Western blot analysis of pAkt protein level in skeletal muscles of *mdx* mice 21 days post Pten inhibitor VO-OHpic treatment. n=4 mice (Control and VO-OHpic). (B) Growth rate of *mdx* mice during VO-OHpic treatment. n=4 mice (Control and VO-OHpic). (C) Muscle weight of *mdx* mice 21 days post VO-OHpic treatment. n=3 mice (Control and VO-OHpic). (D) Representative HE staining image of Qu muscle cross-sections of *mdx* mice 21 days post VO-OHpic treatment. Scale bar: 200 μm for 2.5x, 100 μm for 10x. (E) Quantification analysis of the percentage of necrotic area on cross-section of Qu muscles 21 days post VO-OHpic treatment. n=7 mice (Control), n=8 mice (VO-OHpic). (F) Representative immunofluorescence staining of eMHC and α-Laminin on cross-section of Gas muscles of *mdx* mice 21 days post VO-OHpic treatment. Scale bar: 100 μm. (G) Quantification analysis of eMHC^+^ cell number on cross-section of Gas muscles of *mdx* mice 21 days post VO-OHpic treatment. n=4 mice (Control and VO-OHpic), each dot represents one image at 10x magnification and more than 30 images were taken from 4 mice for each group. Data are means ± s.e.m. Two-tailed *t*-test: \**P* < 0.05; \*\**P* < 0.01, n.s., *P* > 0.05.

